# Investigating variant and expression of CVD genes associated phenotypes among high-risk Heart Failure patients

**DOI:** 10.1101/2023.01.24.525457

**Authors:** Zeeshan Ahmed, Saman Zeeshan, Nicholas Persaud, Bruce T. Liang

## Abstract

Cardiovascular disease (CVD) is a leading cause of premature mortality in the US and the world. CVD comprises of several complex and mostly heritable conditions, which range from myocardial infarction to congenital heart disease. Here, we report our findings from an integrative analysis of gene expression, disease-causing gene variants, and associated phenotypes among CVD populations, with a focus on high-risk Heart Failure (HF) patients. We built a cohort using electronic health records (EHR) of consented patients with available samples, and then performed high-throughput whole-genome and RNA sequencing (RNA-seq) of key genes responsible for HF and other CVD pathologies. We also incorporated a translational aspect to our study by integrating genomics findings with patient medical records. This involved linking ICD-10 codes with our gene expression and variant data to identify associations with HF and other CVDs. Our in-depth gene expression analysis revealed differentially expressed genes associated with HF (41 genes) and other CVDs (23 genes). Furthermore, a variant analysis of whole-genome sequence data of CVD patients identified genes with altered gene expression (FLNA, CST3, LGALS3, and HBA1) with functional and nonfunctional mutations in these genes. Our study highlights the importance of an integrative approach that leverages gene expression, genetic mutations, and clinical data that will allow the prioritization of key driver genes for complex diseases to improve personalized healthcare.

## 1. Introduction

Cardiovascular disease (CVD) is one the leading causes of death in the US and the world [1]. CVD is an ‘umbrella’ term for heart diseases, with primary pathologies including but not limited to heart failure (HF), cardiac arrhythmias, venous thromboembolism, cerebrovascular and peripheral arterial disease, coronary heart disease (CHD), coronary artery disease (CAD), atheromatous vascular disease (AVD), and rheumatic and congenital heart diseases [2]. Several multi-omics state-of-the-art studies of the molecular mechanisms of CVDs [3] involved various data analytic approaches, such as genomics, epigenomics, transcriptomics, proteomics, and metabolomics [3]. Due to the complex nature, progression, and heterogeneity of CVDs, personalized treatments are believed to be critical [4]. Precision medicine is one of the most trending subjects in life and medical science, which promotes intelligently integrative clinical (text and images) and multi-omics data analysis by application of artificial intelligence techniques to provide new insights into CVD [5]. However, we are only in the developing stages of identifying causes of end-stage CVDs [6] and providing better personalized treatments with predictive analysis [7] and deep phenotyping [8].

CVDs are complex and partially heritable conditions, which range from myocardial infarction to CHD [9]. Genomic information, including high-quality sequenced DNA and RNA sequencing (RNA-seq) of transcribed genes, informs us of a CVD patient’s inherent genetic makeup with the most comprehensive view of the genome [10]. We aim to support precision medicine in HF and other CVDs by investigating genes associated with CVDs and analyzing their variants and expression levels that correlate with disease phenotype. RNA-seq-driven gene expression analysis has the potential to reveal novel and sensitive biomarkers and stratify CVD patient populations based on their disease risk [11]. While DNA-based gene variant analysis can assist in diagnosis and plan personalized treatments, integrating and/or validating sequential—one after another—gene expression and variant analyses will certainly advance our understanding of CVD biology. However, several studies have demonstrated that not all genomic variants are detected by transcriptomics as RNA-seq data is only composed of coding regions and cannot detect non-coding mutations [12]. Furthermore, only limited information is available on the completeness of genetic variants detected using RNA-Seq data [13].

Various genomics studies have been conducted recently to discover underlying genetic etiology in CVD patients, especially those suspected of HF disease [14]. Several CVD-related genes have significant mutations and expression differences among CVD patients [15]. In this manuscript, we report a multifaceted approach involving whole-genome sequencing (WGS) integrated with differential gene expression for HF and other CVD study [16, 17].

## 2. Material and Methods

Our research methodology and overall study design (Figure 1) started with analyzing electronic health records (EHR) and building a cohort for CVD sample collection. Next, we implemented experiments by preparing libraries for high-throughput sequencing of extracted DNA and RNA. We then performed WGS and RNA-seq data processing and quality checking, and relational database modelling of extracted variants and genes associated with CVD. We investigated genes responsible for HF and other CVD pathology, and analyzed gene expression, enrichment, annotation, and disease-causing gene variant analysis to identify mutations and changes in mRNA abundance that correlate with HF and CVDs to precisely stratify, classify, and distinguish gender- and age-based patient susceptibility to CVD risks and for genomic predictions of complex phenotypes.

**Figure 1.**
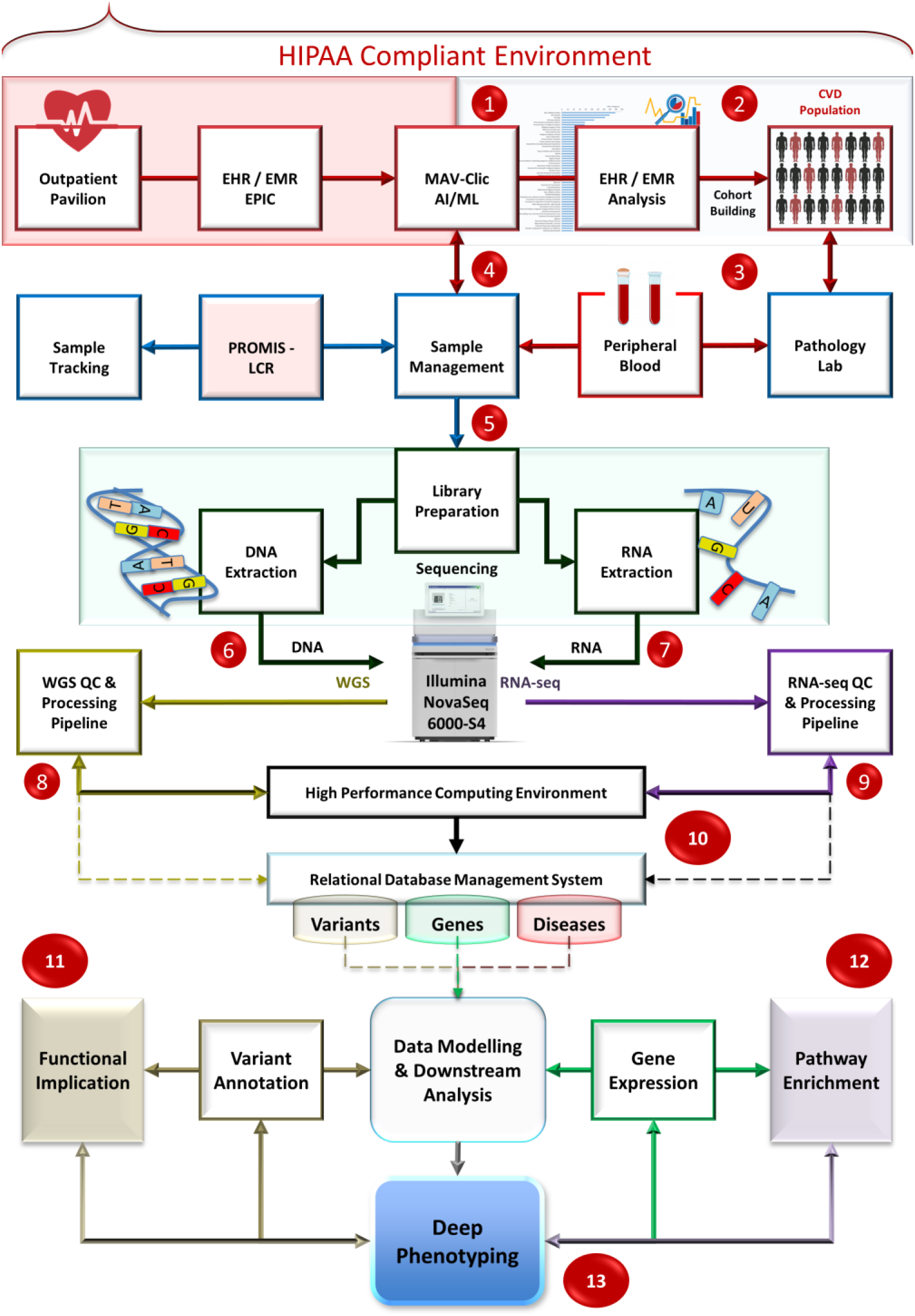
Study design. Overall research methodology is based on the following thirteen steps: 1) clinical data analysis; 2) cohort building; 3) cardiovascular disease (CVD) sample collection; 4) sample management and tracking; 5) Library preparation, and DNA and RNA extraction; 6) high-throughput Whole genome sequencing (WGS); 7) RNA-seq; 8) WGS data processing and quality checking; 9) RNA-seq data processing and quality checking; 10) CVD gene-disease annotation and phenotyping; 11) Gene differential expression and pathway enrichment analysis; 12) Variant and functional mutation analysis; and 13) Deep phenotyping.

### 2.1. Clinical data extraction, CVD cohort building, and high-throughput sequencing

We hypothesize that integrative, intelligent, and analytic access to the EHR has potential to revolutionize the field of medicine by providing the best strategies to diagnose and treat patients at risk for serious medical complications arising from lack of understanding of the biology. However, current limitations imply gaps in clinical and academic settings, difficulties in getting exigent approvals and timeliness of data availability, levels of granularity in clinical information, and application of appropriate modelling strategies that allow learning in the data continuum. Supporting this study, we have developed, tested, and evaluated a HIPAA-compliant intelligent health system named management, analysis, and visualization of clinical data (MAV-clic) [18]. MAV-clic was founded on the scientific premise that to improve the quality and transition of healthcare, integrative platforms are necessary to analyze heterogeneous clinical, epidemiological, metabolomics, and genomics data for precision medicine. It extracts, cleanses, and encrypts clinical data from electronic health systems (EPIC), and then restructures and aggregates them in a deidentified format. Its graphical user interface allows user-friendly interactive querying, analysis, and visualization of clinical data. MAV-clic is based on product-line architecture, which implies that all major designed and developed modules of the system can perform individual key roles but are integrated with each other as well as [18].

To support this study, we also developed an efficient data management system (PROMIS-LCR) for patient recruitment and consent, and for collecting, storing, and tracking of the original and current quantities of biospecimens under standardized conditions for preservation of critical metabolites [19]. PROMIS-LCR has been successfully tested, evaluated, and deployed, and is operational at the UConn Outpatient Pavilion (OP) to support establishment of a biobank and a precision medicine study at UConn Health. MAV-clic is fully integrated with PROMIS-LCR, and operational to efficiently extract and link de-identified medical details of the consented patients participating in this study with their collected biospecimens. With integrated MAV-clic and PROMIS-LCR, we analyzed patient population data to build a CVD cohort centered on medical details, symptoms, age, race, gender, and demographics. We extracted DNA from 61 CVD patients: 40 male and 21 female individuals (65.57% male and 34.42% female population) aged between 45 to 92, with diverse ethnic groups (56 Not Hispanic, 4 Hispanic, and 1 Declined to Answer), and self-described race (42 Whites, 7 Blacks or African Americans, 1 Asian, and 11 of unknown race). These patients were clinically diagnosed with CVD and CMS/HCC HF, as well as cardiomyopathy, hypertension, obesity, type 2 diabetes mellitus, asthma, high cholesterol, hernia, chronic kidney, joint pain, myalgia, dizziness and giddiness, osteopenia of multiple sites, chest pain, and osteoarthritis. To support this study, we also extracted DNA from 10 healthy individuals (5 males and 5 females; out of which 3 were self-described Hispanics and 7 non-Hispanic; 9 were White race and 1 unknown race) aged between 28 to 78 with no clinical manifestation of CVD.

We created a Next-generation sequencing (NGS) CVD dataset, which includes high-quality sequenced (WGS and RNA-seq, Illumina NovaSeq6000-S4) data of randomly consented 61 patients, available with their personal medical history. Paired-end 150 bp short sequences (reads, pool across 2 lanes) with 30X coverage were generated from DNA and RNA isolated from peripheral blood samples, including the Illumina-compatible library (TruSeq Stranded mRNA). RNA quality was assessed for RNA integrity number, which was >7 for all samples. Written informed consent was obtained from all subjects. All procedures performed in studies involving human participants were in accordance with the ethical standards of the institution and with the 1964 Helsinki declaration and its later amendments or comparable ethical standards. All human samples were used in accordance with relevant guidelines and regulations, and all experimental protocols were approved by the Institutional Review Board (IRB) at UConn Health.

### 2.2. WGS data processing, quality checking, analysis, and visualization

We applied our in-house developed gene-variant analysis pipeline (JWES) for the whole genome data pre-processing, modelling, and downstream analysis. JWES processes the raw sequence data, converts raw signals into base calling, identifies regions of interest in the genome, aligns, assembles contigs and scaffolds, and detects variants [20]. Its operations are divided into three separate modules: I) data preprocessing, II) storage and management, and III) visualization. JWES is a cross-platform java-based application that integrates multiple open-source command-line tools for sequence data processing and analysis [20].

We analyzed the mutations in genes implicated in CVDs and HF as coding proteins with the labeled recurrent hotspots. The variant data were annotated for biological and functional implications with three computational algorithms: Scale-Invariant Feature Transform (SIFT) [21, 22, 23], Polymorphism Phenotyping v2 (PolyPhen-2) [24], and MutationAssessor [25, 26]. We used SIFT to predict the effect of coding variants on protein function [22], as the challenge here is to identify causative variants for the phenotypes linked to HF; SIFT was originally proposed to predict the impact of amino acid substitutions on protein function [26]. PolyPhen-2 is an automatic web tool we used for the extraction of sequences and structure-based features of the substitution site. The SNPs were analyzed in a batch mode, predicted for the functional impact, and searched for in a database of precomputed predictions for the WGS data. We applied MutationAssessor to differentiate between conserved patterns using conservation and specificity score to account for functional shifts between subfamilies of proteins [56]. MutationMapper [27,28] was used to generate the lollipop plots for all the genes (Figures 4–6). These plots display the highly recurrent mutations (nucleotide alterations) including mutations with low frequency.

**Figure 2.**
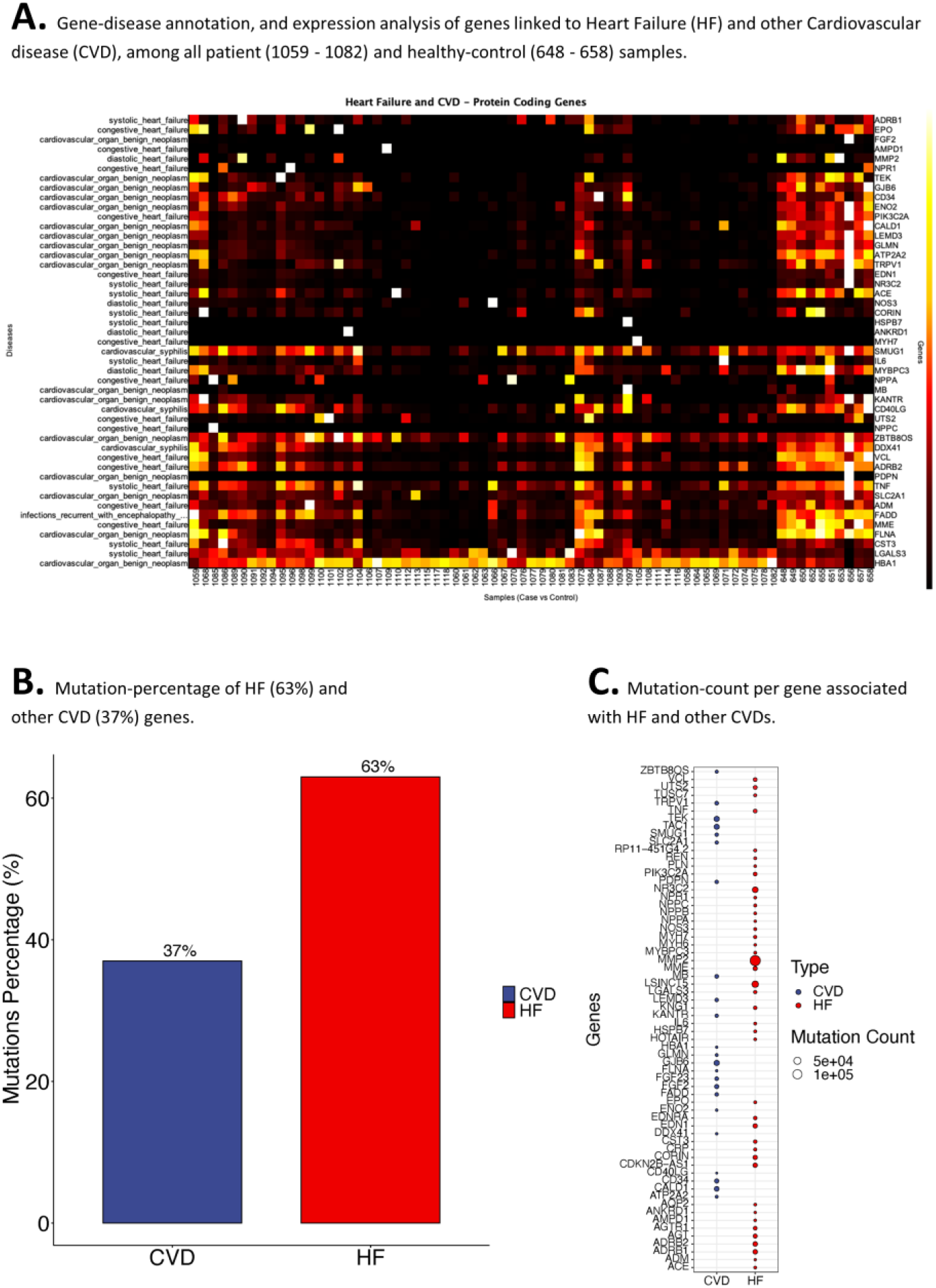
Gene-disease annotation, expression and variant analysis of genes associated with Heart Failure (HF) and other cardiovascular diseases (CVDs). HF genes include, TNF, IL6, ACE, MMP2, NOS3, AGT, EDN1, REN, MYH7, AGTR1, NPPA, ADRB2, NR3C2, MME, CRP, MYH6, EPO, CST3, EDNRA, AQP2, MYBPC3, KNG1, VCL, HOTAIR, CDKN2B-AS1, ANKRD1, ADM, AMPD1, PLN, LGALS3, NPPB, ADRB1, UTS2, PIK3C2A, NPPC, CORIN, NPR1, LSINCT5, TUSC7, HSPB7, and RP11-451G4.2. Other CVD genes include, SLC2A1, FGF2, FLNA, HBA1, GJB6, ATP2A2, CD40LG, FGF23, TEK, TAC1, DDX41, FADD, ENO2, LEMD3, CD34, TRPV1, GLMN, MB, SMUG1, PDPN, CALD1, KANTR, and ZBTB8OS. Figure 3A) Gene-disease annotation, phenotyping, and expression analysis of protein-coding genes associated with heart failure (HF) and other cardiovascular diseases (CVD). Figure 3B) Mutation-percentage of HF (63%) and other CVD (37%) genes. Figure 3C) Mutation-count per gene associated with HF and other CVDs.

**Figure 3.**
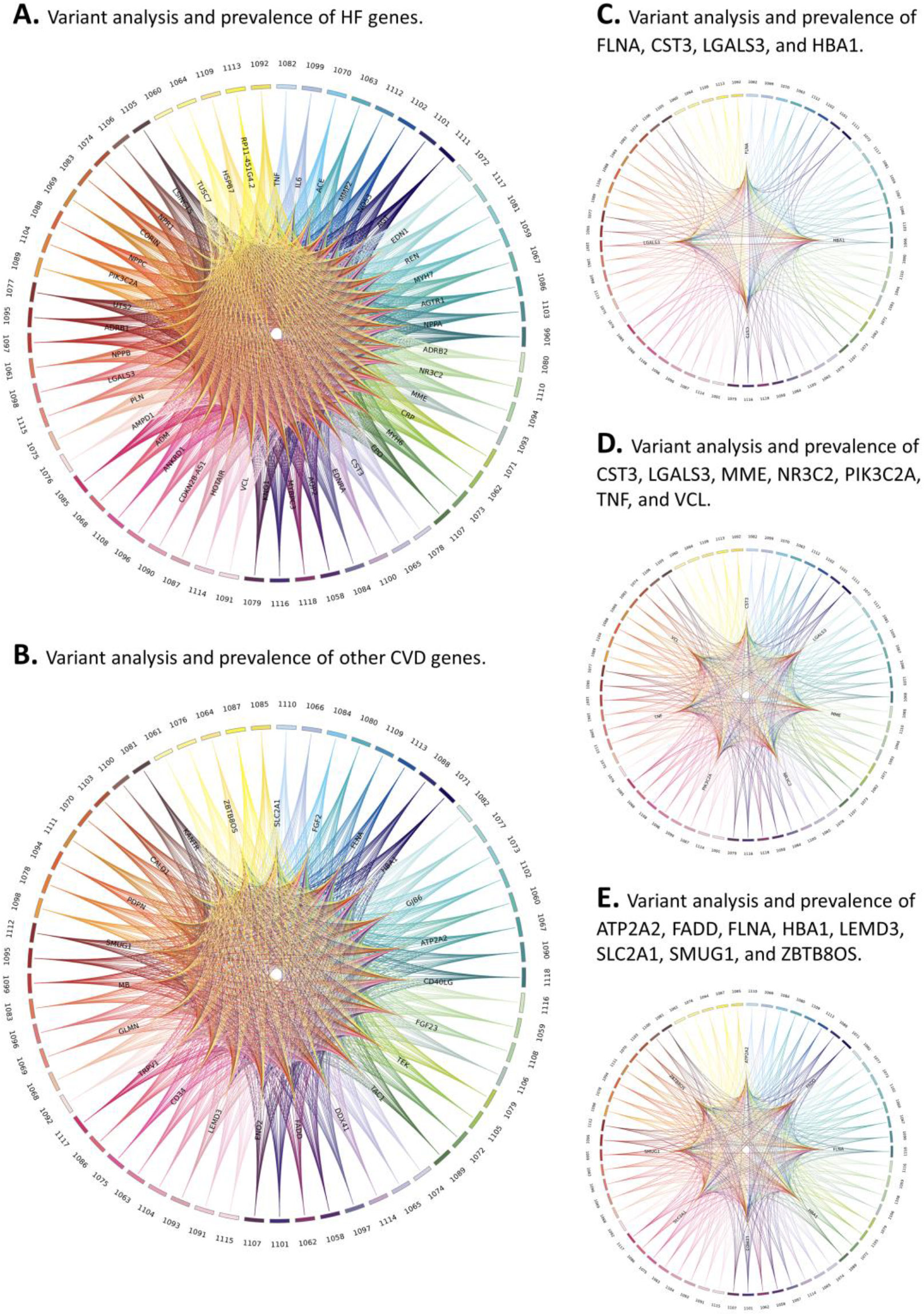
Gene-variant analysis of protein-coding genes associated with heart failure (HF) and other cardiovascular diseases (CVD). Figure 3A) Variant analysis and prevalence of HF genes. Figure 3B) Variant analysis and prevalence of other CVD genes. Figure 3C) Variant analysis and prevalence of FLNA, CST3, LGALS3, and HBA1. Figure 3D) Variant analysis and prevalence of CST3, LGALS3, MME, NR3C2, PIK3C2A, TNF, and VCL. Figure 3E) Variant analysis and prevalence of ATP2A2, FADD, FLNA, HBA1, LEMD3, SLC2A1, SMUG1, and ZBTB8OS.

**Figure 4.**
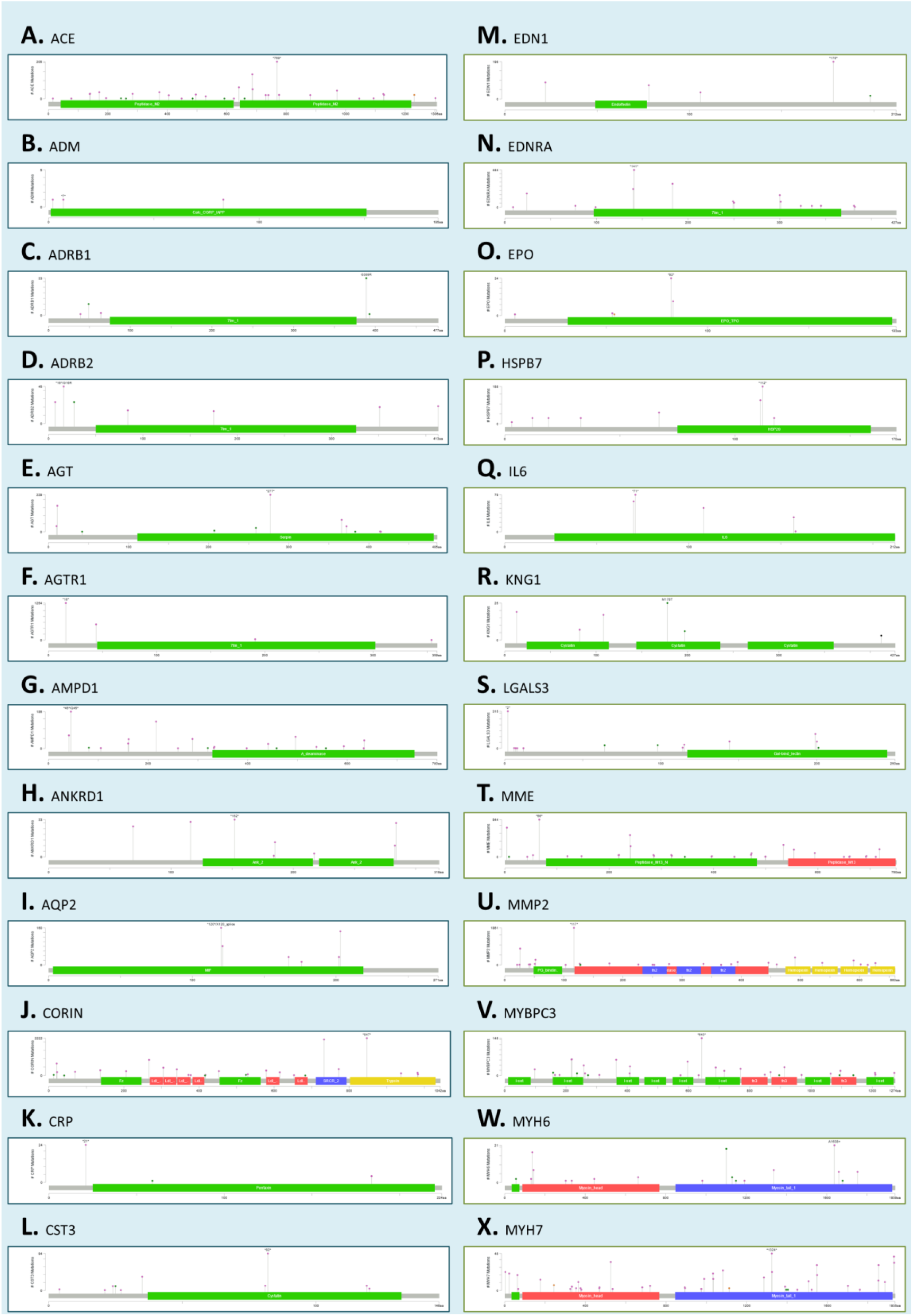
Functional and non-functional mutation analysis of genes associated with heart failure. Lollipop plots of ACE, ADM, ADRB1, ADRB2, AGT, AGTR1, AMPD1, ANKRD1, AQP2, CORIN, CRP, CST3, EDN1, EDNRA, EPO, HSPB7, IL6, KNG1, LGALS3, MME, MMP2, MYBPC3, MYH6, and MYH7. Green color represents Missense Mutations, black represent Truncating Mutations, brown represent Inframe Mutations, and purple represent Fusion Mutations.

**Figure 5.**
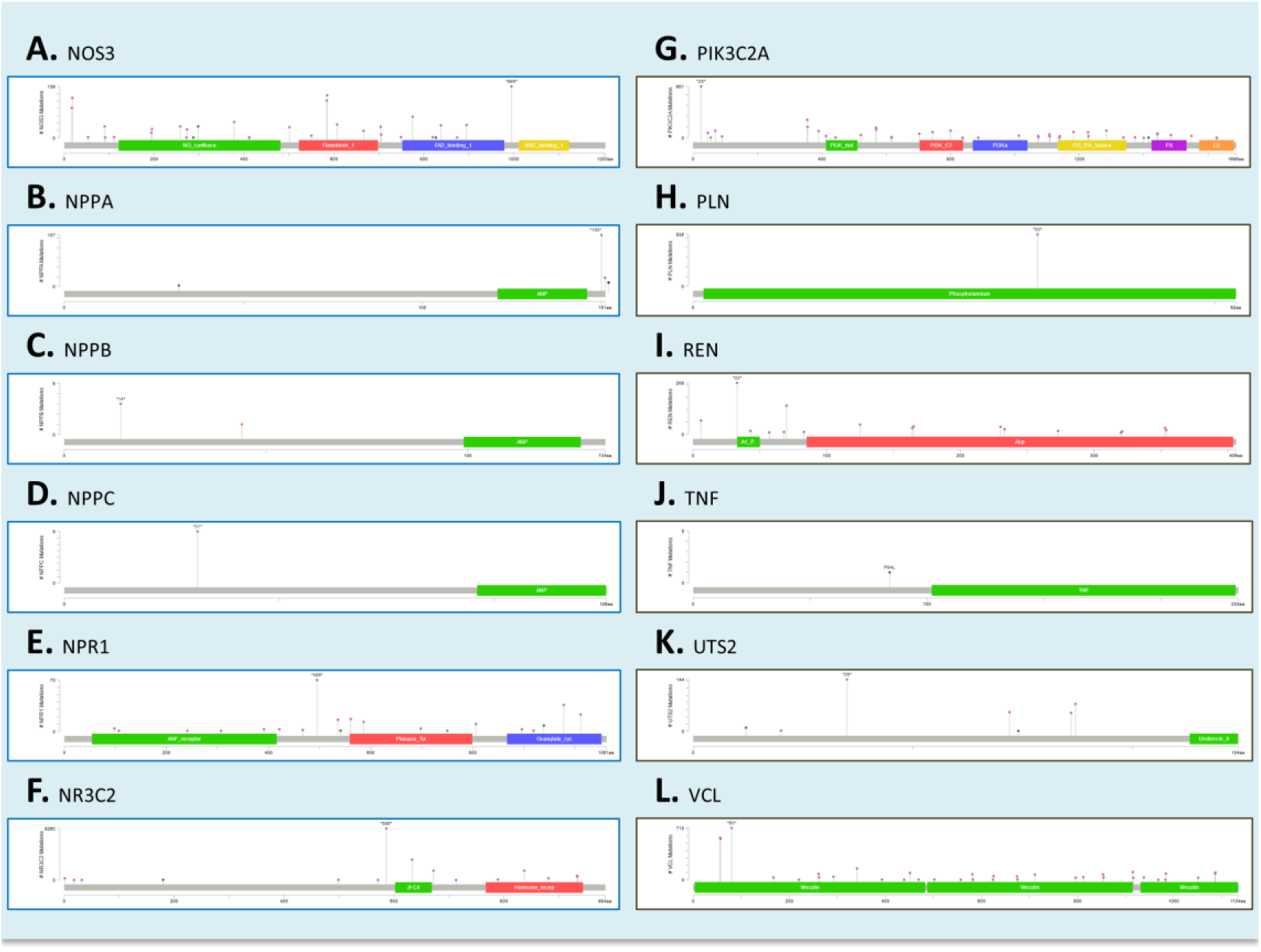
Functional and non-functional mutation analysis of genes associated with heart failure. Lollipop plots of NOS3, NPPA, NPPB, NPPC, NPR1, NR3C2, PIK3C2A, PLN, REN, TNF, UTS2, and VCL. Green color represents Missense Mutations, black represent Truncating Mutations, brown represent Inframe Mutations, and purple represent Fusion Mutations.

**Figure 6.**
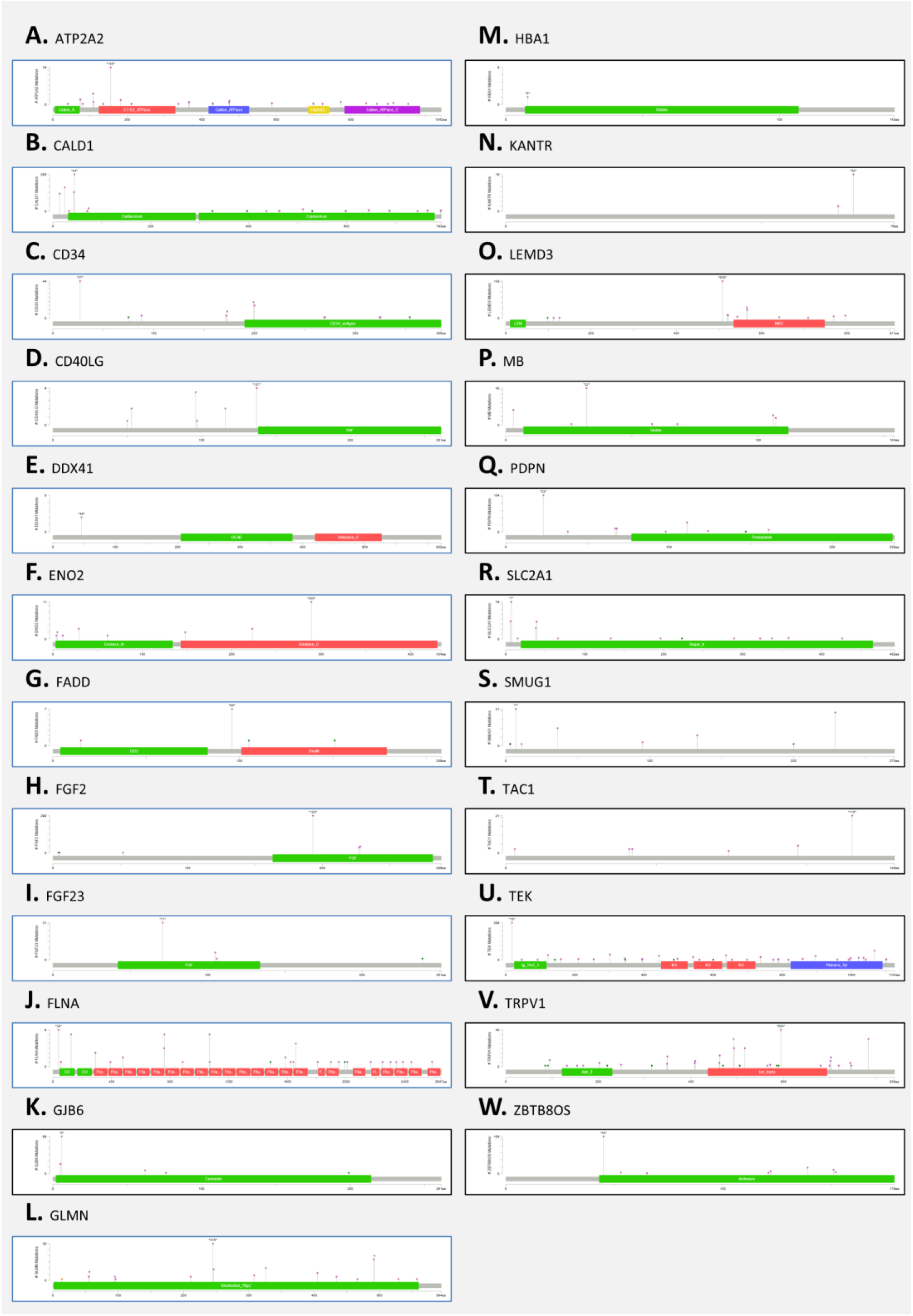
Functional and non-functional mutation analysis of genes associated with other cardiovascular disease (CVDs). Lollipop plots of SLC2A1, FGF2, FLNA, HBA1, GJB6, ATP2A2, CD40LG, FGF23, TEK, TAC1, DDX41, FADD, ENO2, LEMD3, CD34, TRPV1, GLMN, MB, SMUG1, PDPN, CALD1, KANTR, and ZBTB8OS. Green color represents Missense Mutations, black represent Truncating Mutations, brown represent Inframe Mutations, and purple represent Fusion Mutations.

### 2.3. RNA-seq data processing, quality checking, analysis, and visualization

We developed a detailed bioinformatics pipeline for RNA-seq data pre-processing, and gene expression data analysis, annotation with relevant diseases, and heatmap visualization without requiring a strong computational background from the user [17]. RNA by expectation maximization (RSEM) [29] is applied for the quantification and identification of differentially expressed genes (DEGs) and aligns sequences to reference de novo transcriptome assemblies [29]. Results of the processed RNA-seq data are automatically parsed and uploaded into our designed relational database. The outcome of the pipeline includes but is not limited to the quality metrics, and quantified gene and isoform abundances with transcripts per million (TPM), fragments per kilobase million (FPKM), reads per kilobase of transcript per million mapped reads (RPKM), and mean expressed transcript lengths. TPM values are proven to be more accurate measures of true abundance of RNA molecules from given genes and are more consistent across libraries. This allows a more stable statistical analysis. Case-control studies for gene expression analysis also involved Gene Set Enrichment Analysis (GSEA) [30] to associate cellular functions with the DSGs and verify the differences between comparisons. Following the standard procedure described by the GSEA user guide, during GSEA, we utilized Molecular Signature Database v7.0 gene lists of hallmark gene sets (H), the Kyoto Encyclopedia of Genes and Genomes (KEGG) pathway database (C2), and the REACTOME pathway database (C2).

We interfaced the RNA-seq pipeline with our newly developed robust bioinformatics, open-source, user-friendly, cross-platform, desktop, and database application, i.e., GVViZ (visualizing genes with disease causing variants) [17]. GVViZ follows the FAIR paradigm, based on a set of simple instructions that allow users without computational experience (e.g., bench scientists, non-computational biologists, and geneticists) to analyze, visualize, and export data to share results. It allows the user to 1) select among genes of interest, expression cutoffs, and samples; 2) search and select genes and associated diseases for the analysis; 3) annotate and perform gene expression analysis based on selected abundances and filtering conditions, samples, genes, and diseases; 4) produce dynamic and customized heatmap visualizations; and 5) export results in image and text (CSV) files. To advance data analytic capabilities of GVViZ, we have modelled and implemented an annotated disease-gene-variants knowledgebase based on clinical, genomics, and mutation data collected from several databases and open archives worldwide [31, 32]. These include ClinVar, GeneCards, MalaCard, DISEASES, HGMD, Disease Ontology, DiseaseEnhancer, DisGeNET, eDGAR, GTR, OMIM, miR2Disease, DNetDB, The Cancer Genome Atlas, International Cancer Genome Consortium, OMIM, GTR, CNVD, Ensembl, GenCode, Novoseek, Swiss-Prot, LncRNADisease, Orphanet, WHO, FDA, Catalogue Of Somatic Mutations In Cancer (COSMIC), and Genome-wide Association Studies (GWAS) [64]. Overall, the annotation database includes 59,293 total genes (19,989 protein-coding and 39,304 non-protein-coding genes) and over 200,000 gene-disease combinations [64].

## 3. Results

### 3.1. Gene-disease annotation and expression analysis

In our initial analysis, we integrated our disease-gene-variant database to identify 41 HF and 23 other CVD-related genes (Figure 2A) [16]. Next, we performed gene expression analysis to identify differentially expressed protein coding genes [35]. We identified 4,712 differentially expressed genes (DEGs) in CVD patients. 42 genes were found to be statistically significant with P value < 0.05 and ļlog2FCļ>= 2 showing greater than 2-fold change. Some of these highly significant genes have already been reported (*APOD, PIGR, CELSR1, COBLL1, FCRL5, TEAD2, ABCA6, COL4A3, CYP4F2, FMOD, GNG8, IGF2R, PEG10, RAPGEF3, RASGRF1, SCARNA17, TCF4*), while some genes (*ADAM29, ARHGAP44, CD200, CLEC17A, CLNK, CNTNAP1, CNTNAP2, CTC-454I21.3, DMD, FAM129C, FAM3C, FCRL1, FCRL2, FCRLA, GPM6A, KLHL14, MTRNR2L3, NPIPB5, OSBPL10, PAX5, PCDH9, PHYHD1, POU2AF1, RALGPS2, ZNF888*) have novel expression in CVD. Gene enrichment of the DEGs revealed multiple pathways upregulated (190) and downregulated (408) based on the normalized enrichment score (NES) in the CVD patients [16].

### 3.2. Gene-variant analysis of HF and other CVD genes

We conducted high depth WGS of similar individuals recruited for CVD study [35], we performed variant analysis and verified mutations among the annotated genes for HF (Figure 3A) and CVD patients (Figure 3B), and identified mutations among four genes with altered expression (FLNA, CST3, LGALS3, and HBA1) (Figure 3C). Notables are the missense mutations in FLNA, CT3, and LGALS3. Furthermore, we found mutations in HF genes (ACE, ADRB1, ADRB2, AGT, AMPD1, CORIN, EDN1, KNG1, MME, MYBPC3, MYH6, MYH7, NOS3, NPPA, NPR1, NR3C2, PIK3C2A) (Figure 3D) and other CVD genes (CALD1, CD34, FADD, FLNA, TEK, and TRPV1) that were also differentially expressed (Figure 3E). We annotated these mutations to identify functional and nonfunctional mutations in their sequences of all genes associated with HF and other CVDs.

In total, we detected 1,039,750 single nucleotide variants (SNV) and insertion and deletion events. The most common mutation types in HF and CVD genes were intronic, 5’ flank, and 3’ flank mutations (Figures 4, 5, and 6). Mutations in these genes have been linked to aberrant expression in CVD. Furthermore, the missense mutations common amongst these genes were benign or had low functional impact. We were unable to find or report functional mutations among five genes related to HF: *CDKN2B-AS1* [33]*, HOTAIR* [34]*, LSINCT5* [35], *RP11-451G4.2, and TUSC7* [36]. We could not find functional impact scores of mutations found in most of the genes investigated. Possible deleterious mutations were only found in AGT, AMPD1, LGALS3, and MYBPC3. Further details are provided in Table 1 and Table 2. Detailed functional and non-functional mutation analysis of HF and other CVD genes is attached in the supplementary material.

**Table 1.**
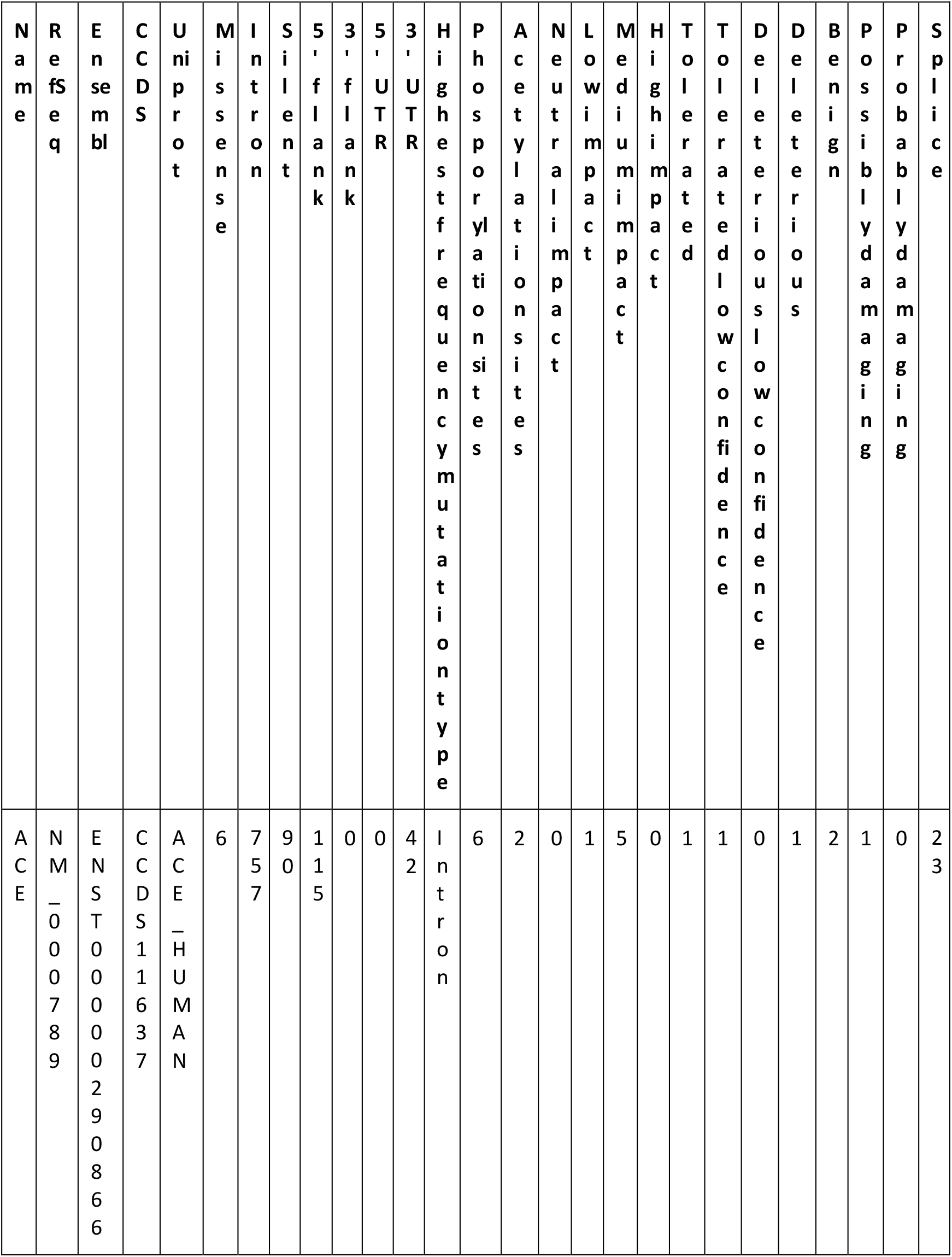
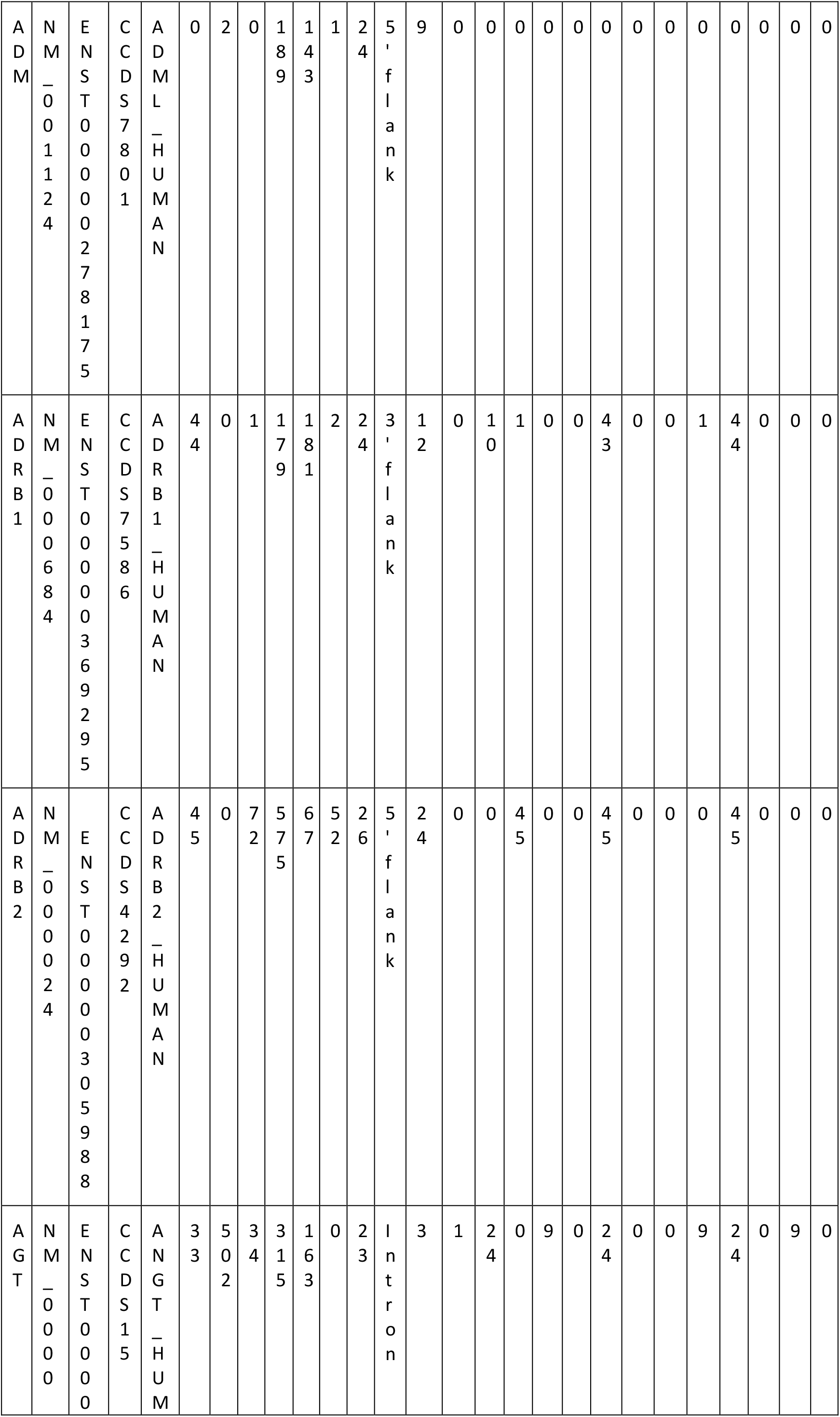
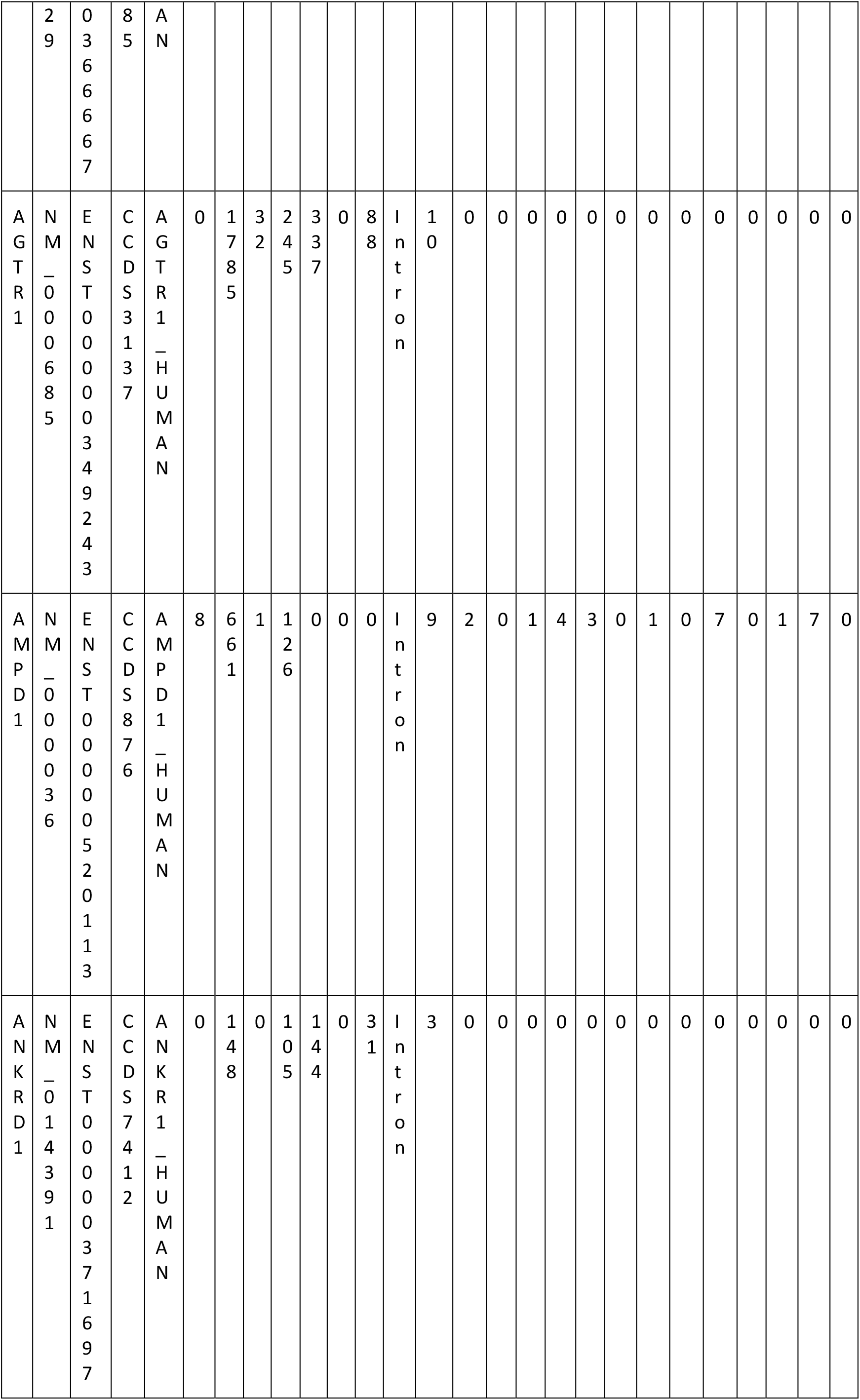
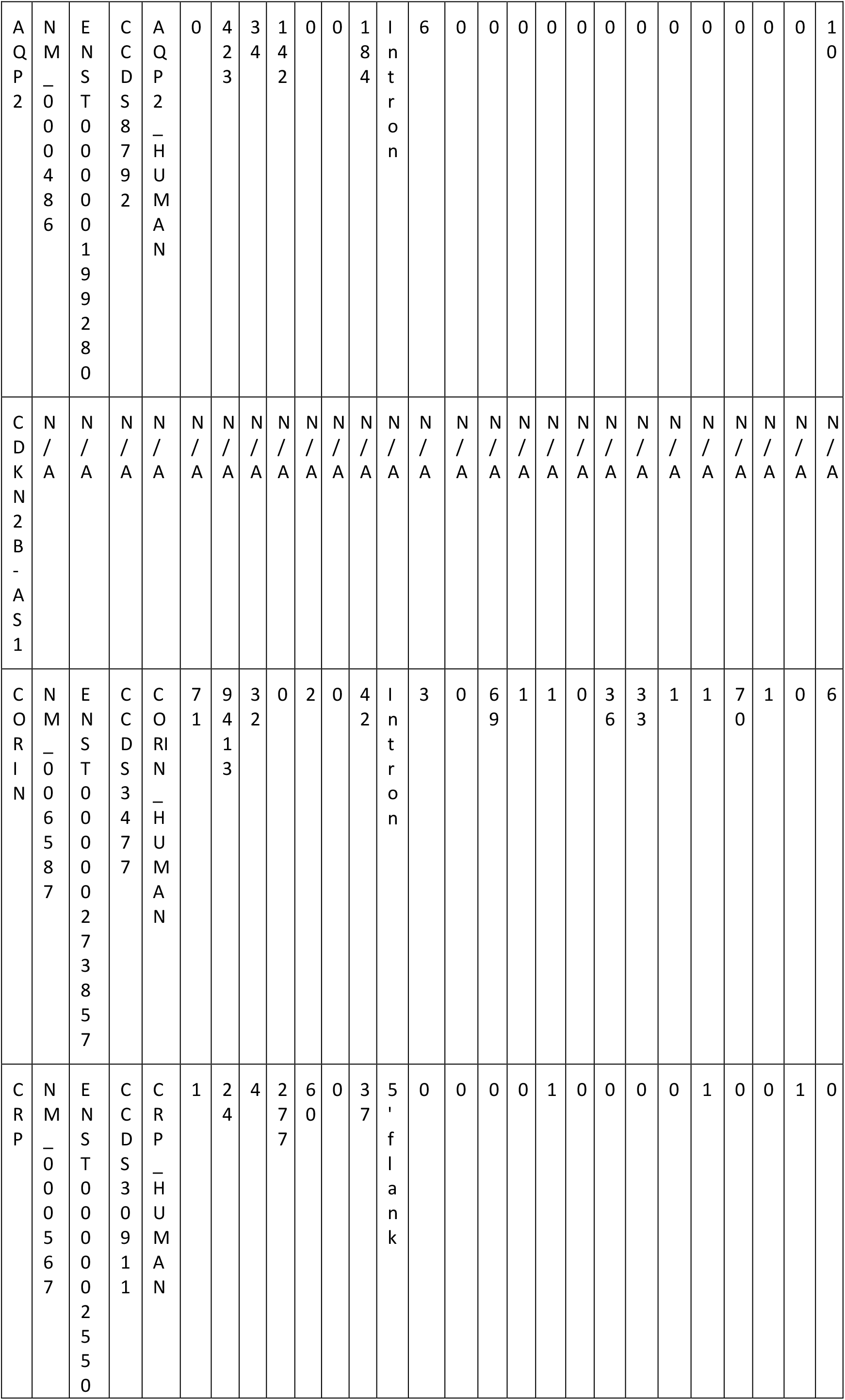
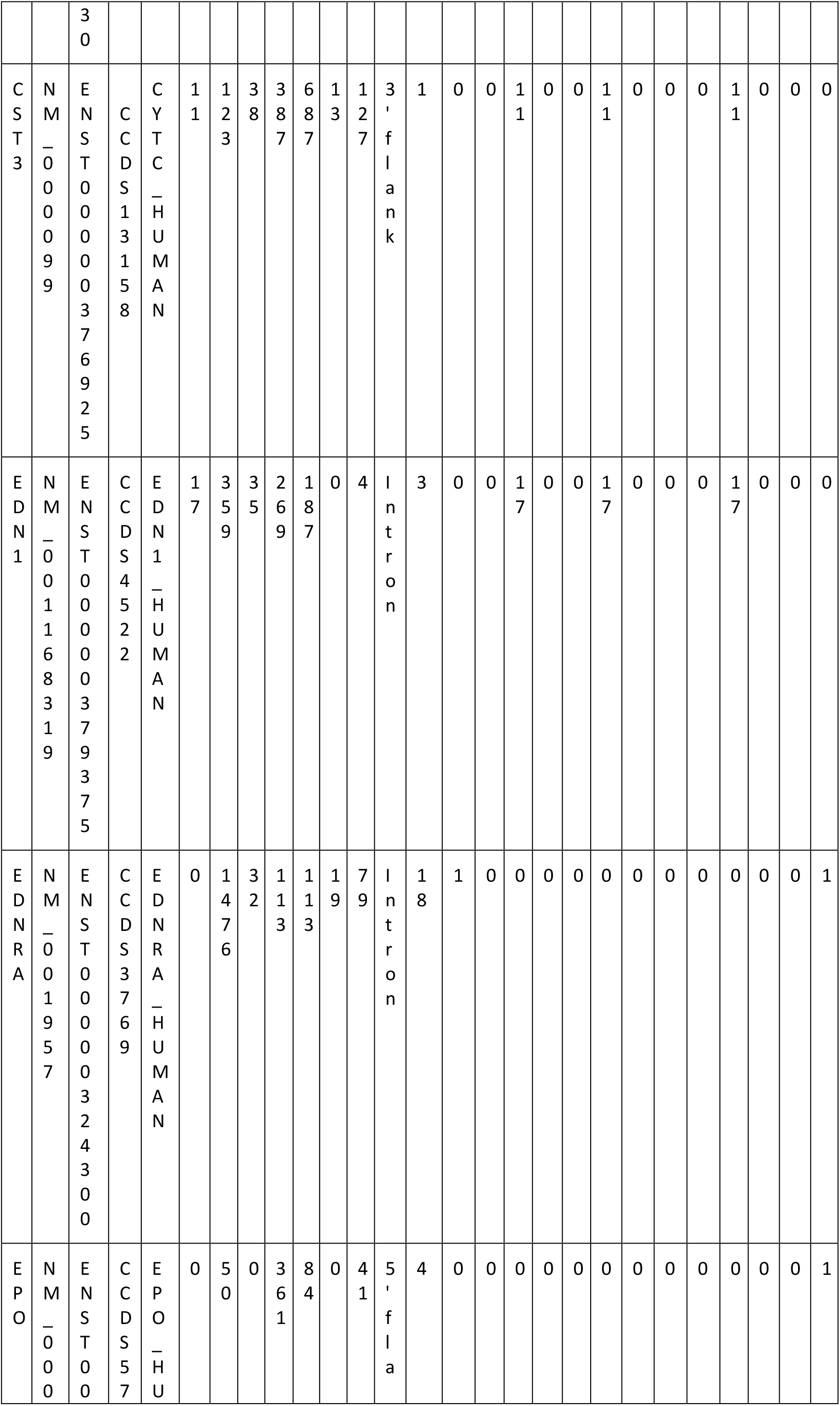
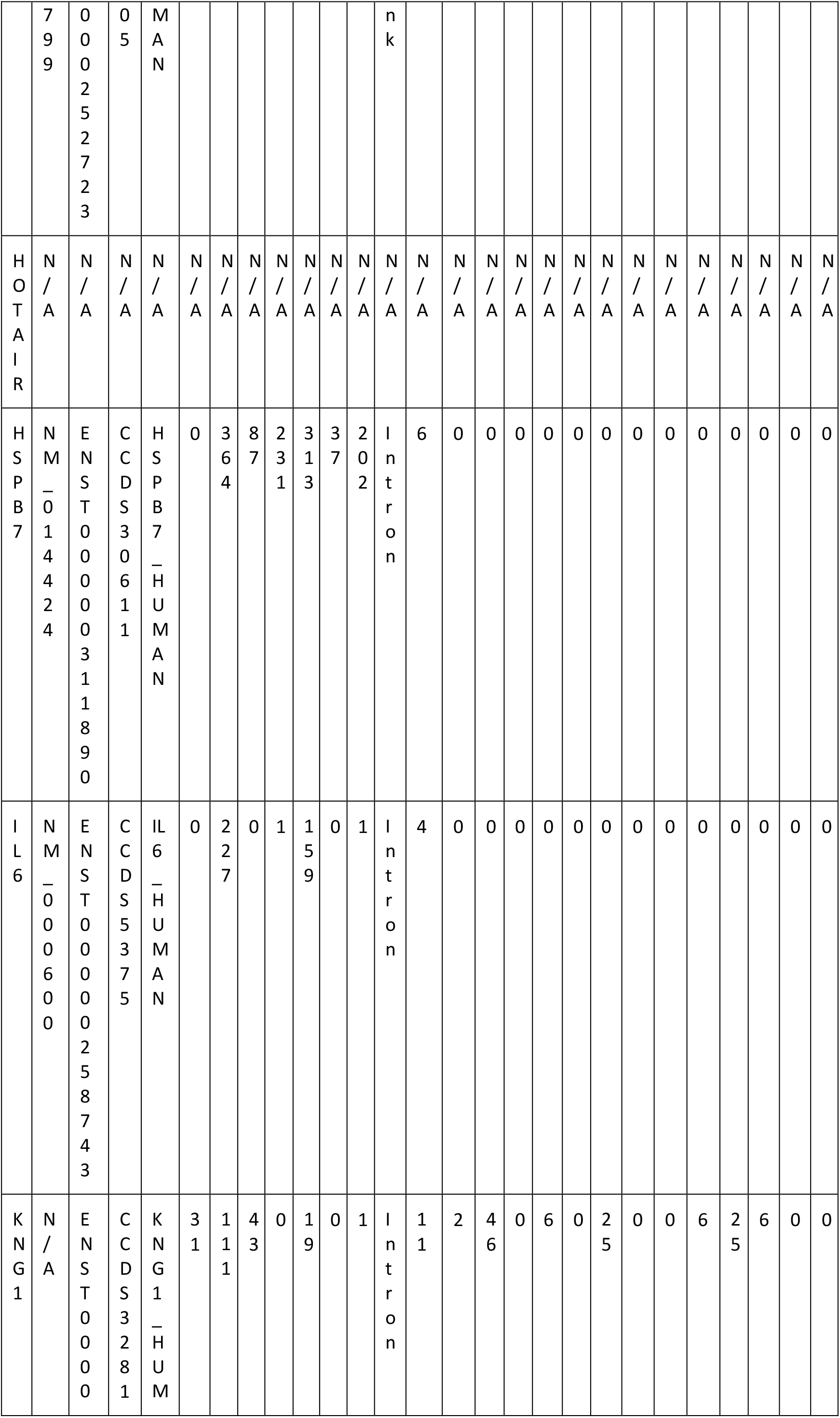
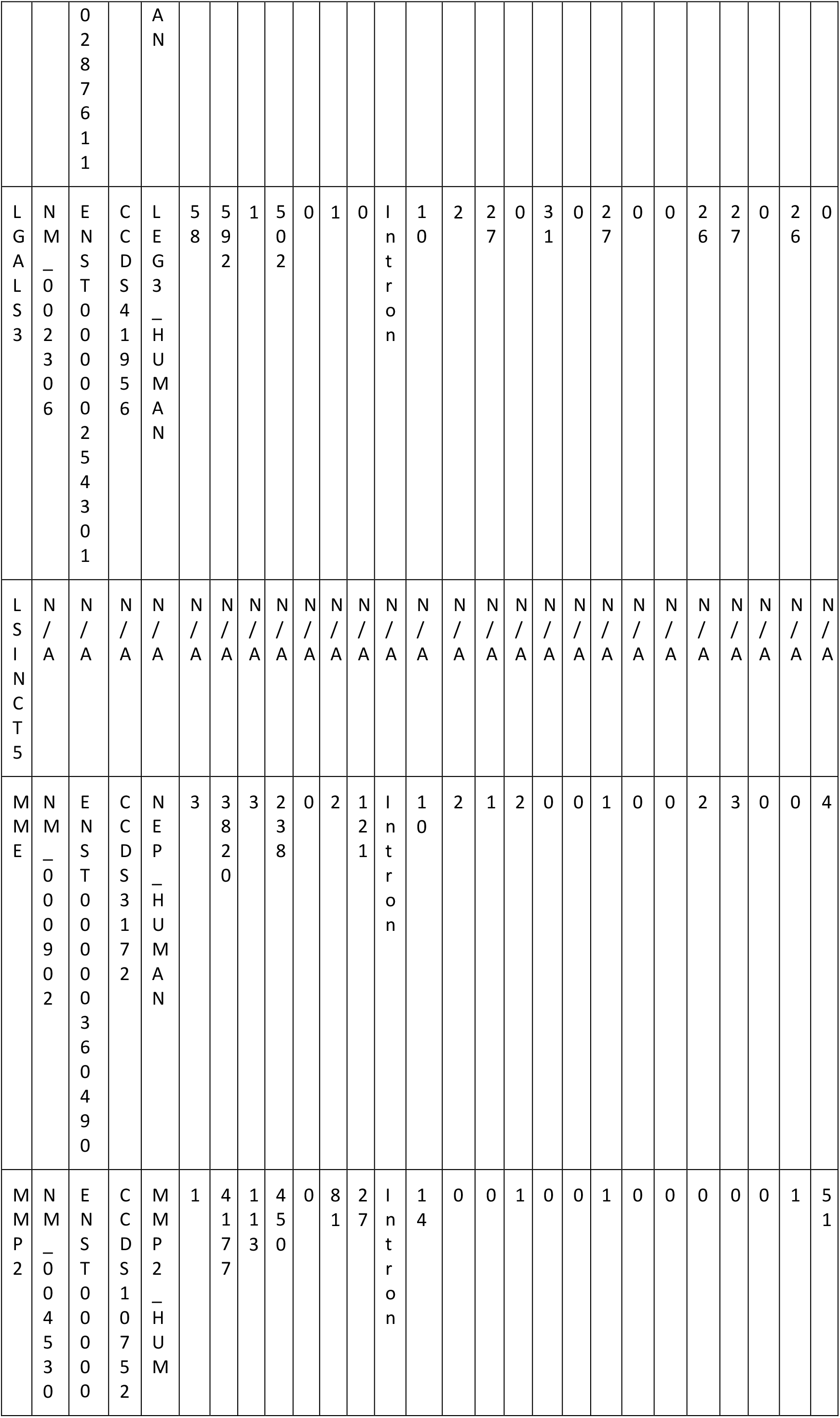
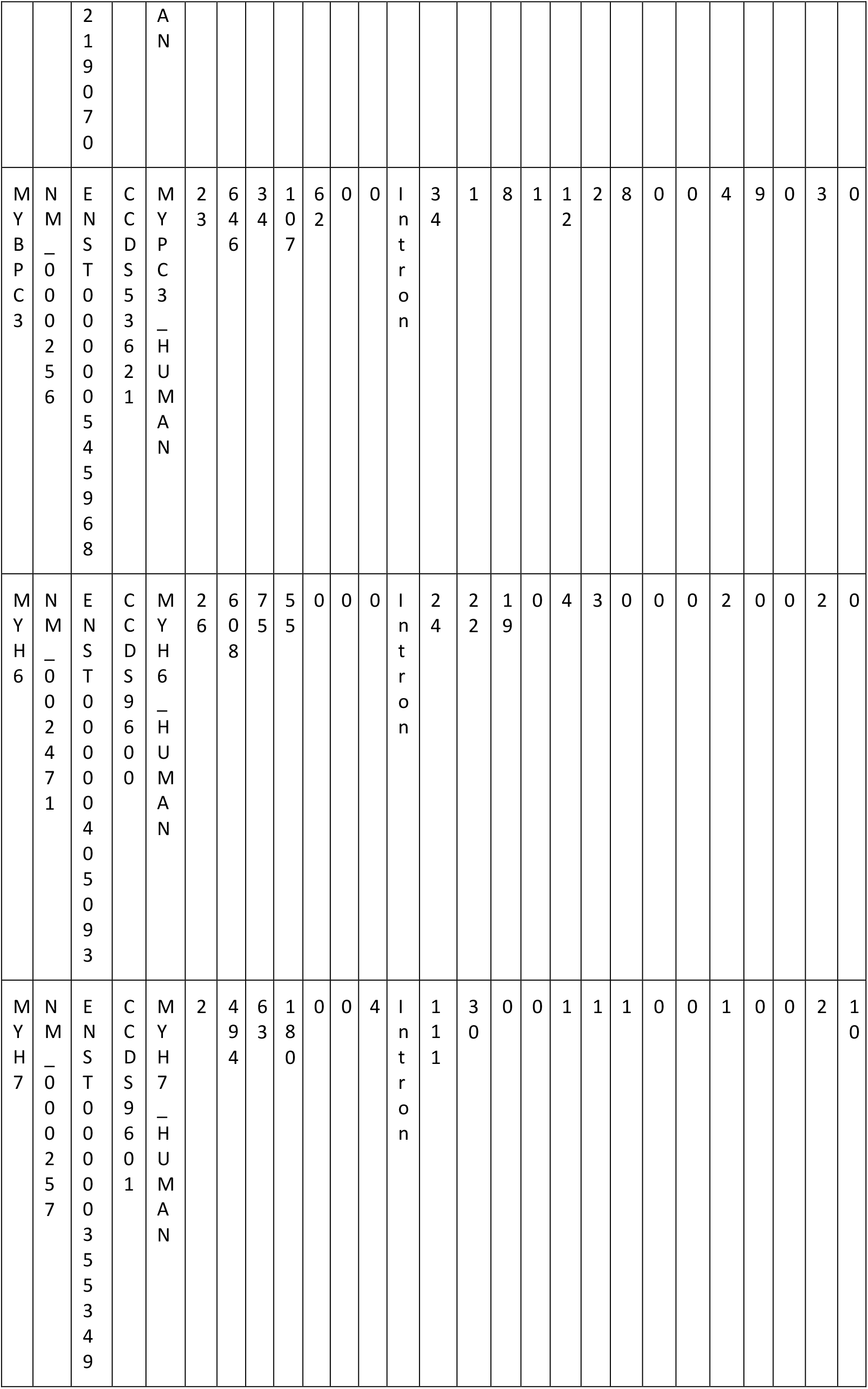
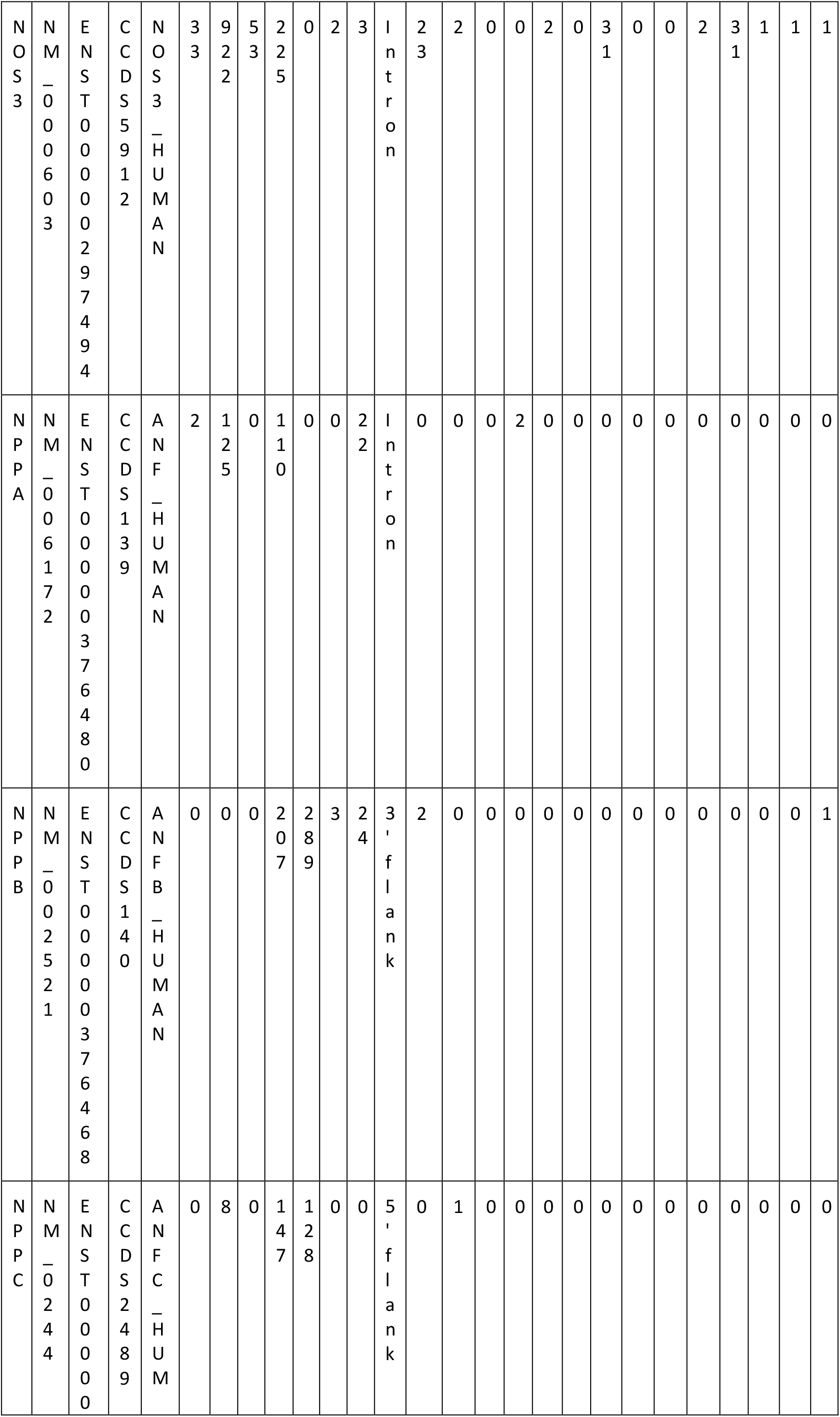
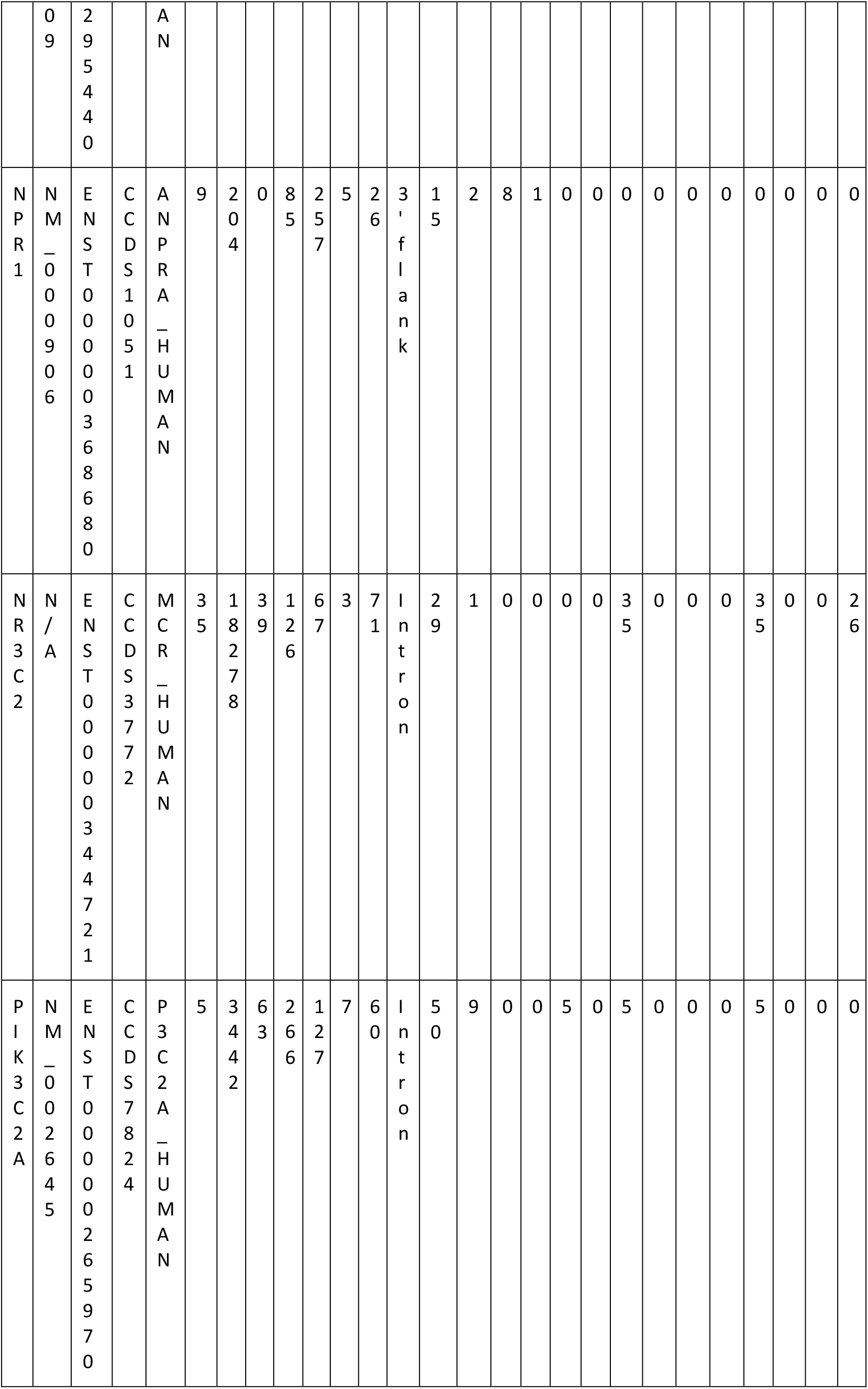
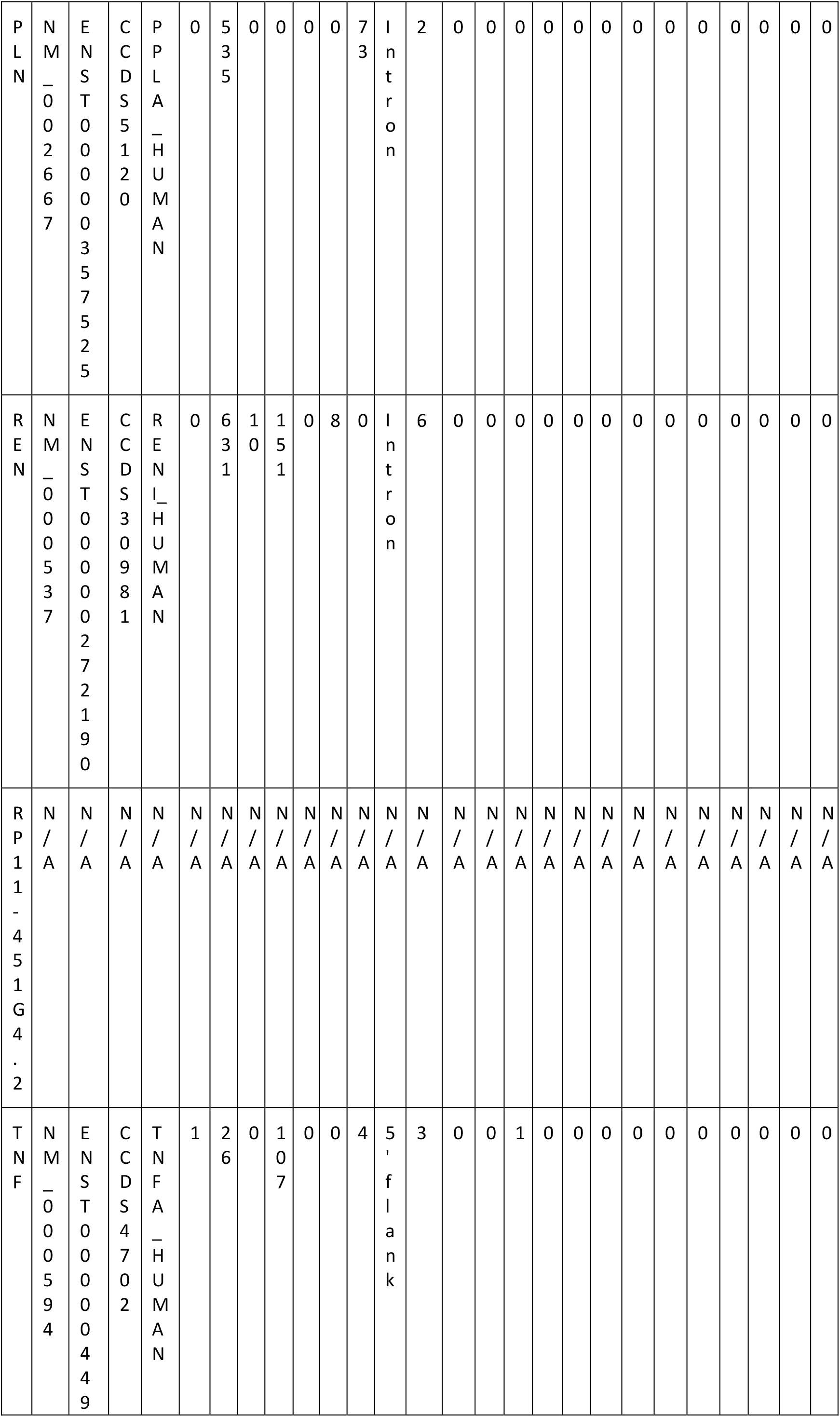
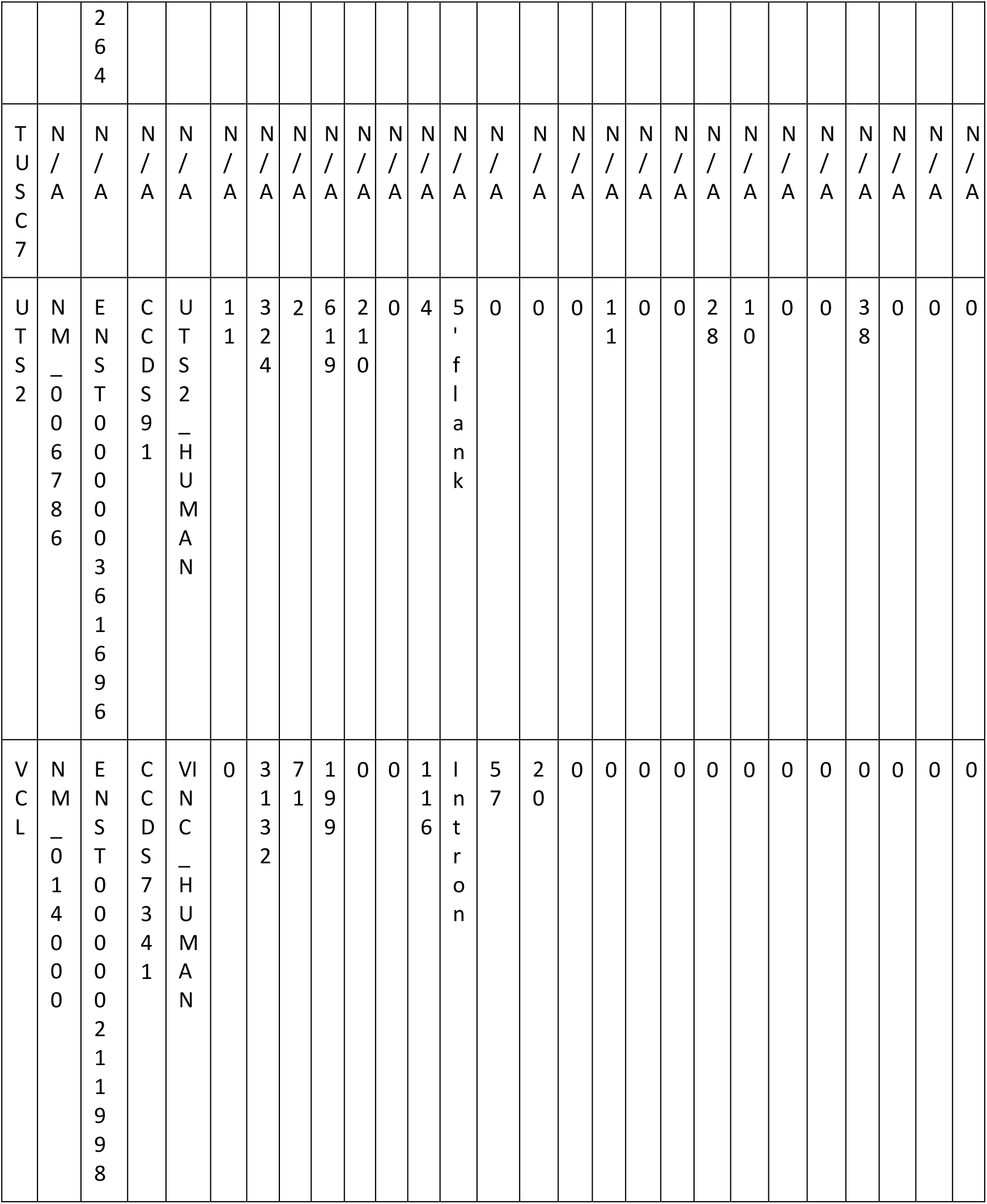
Splice mutation analysis of genes associated with heart failure. Tabulated information includes Name, RefSeq Ensembl, CCDS, Uniprot Missense, Intron, Silent, 5’ flank, 3’ flank, 5’ UTR, 3’ UTR, Highest frequency mutation type, Phosphorylation sites, Acetylation sites, Neutral impact, Low impact, Medium impact, High impact, Tolerated, Tolerated low confidence, Deleterious low confidence, Deleterious, Benign, Possibly damaging, and Probably damaging.

**Table 2.**
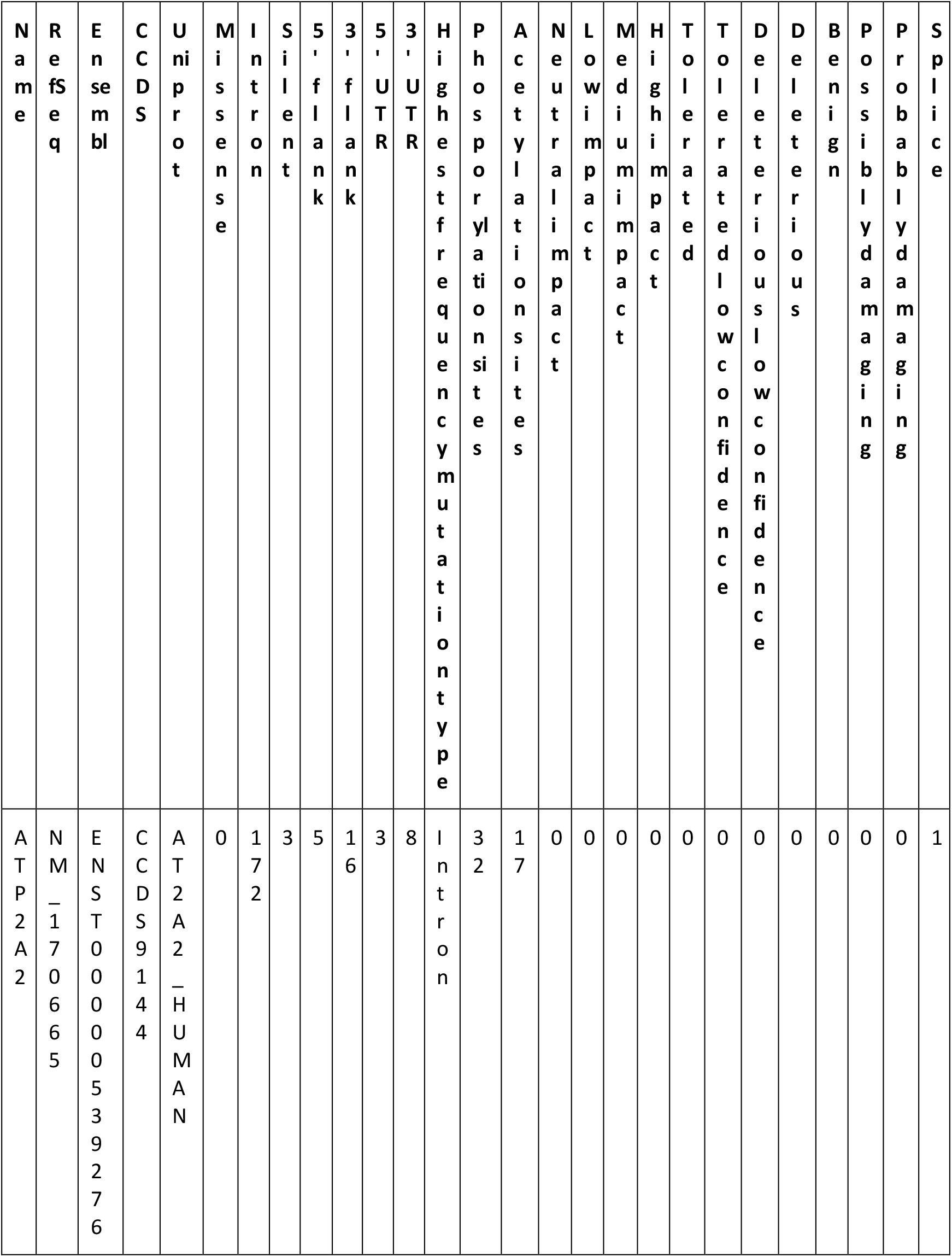
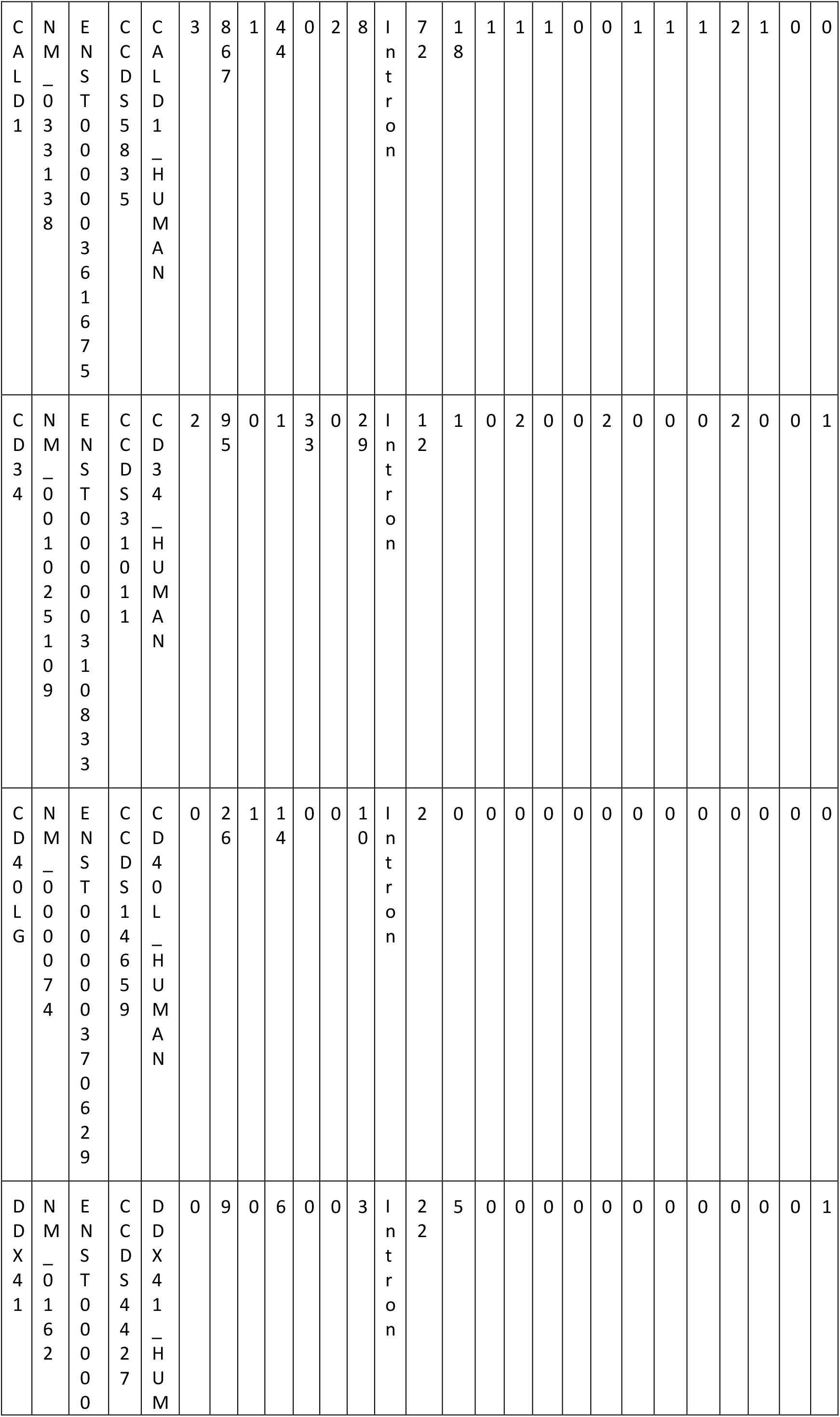
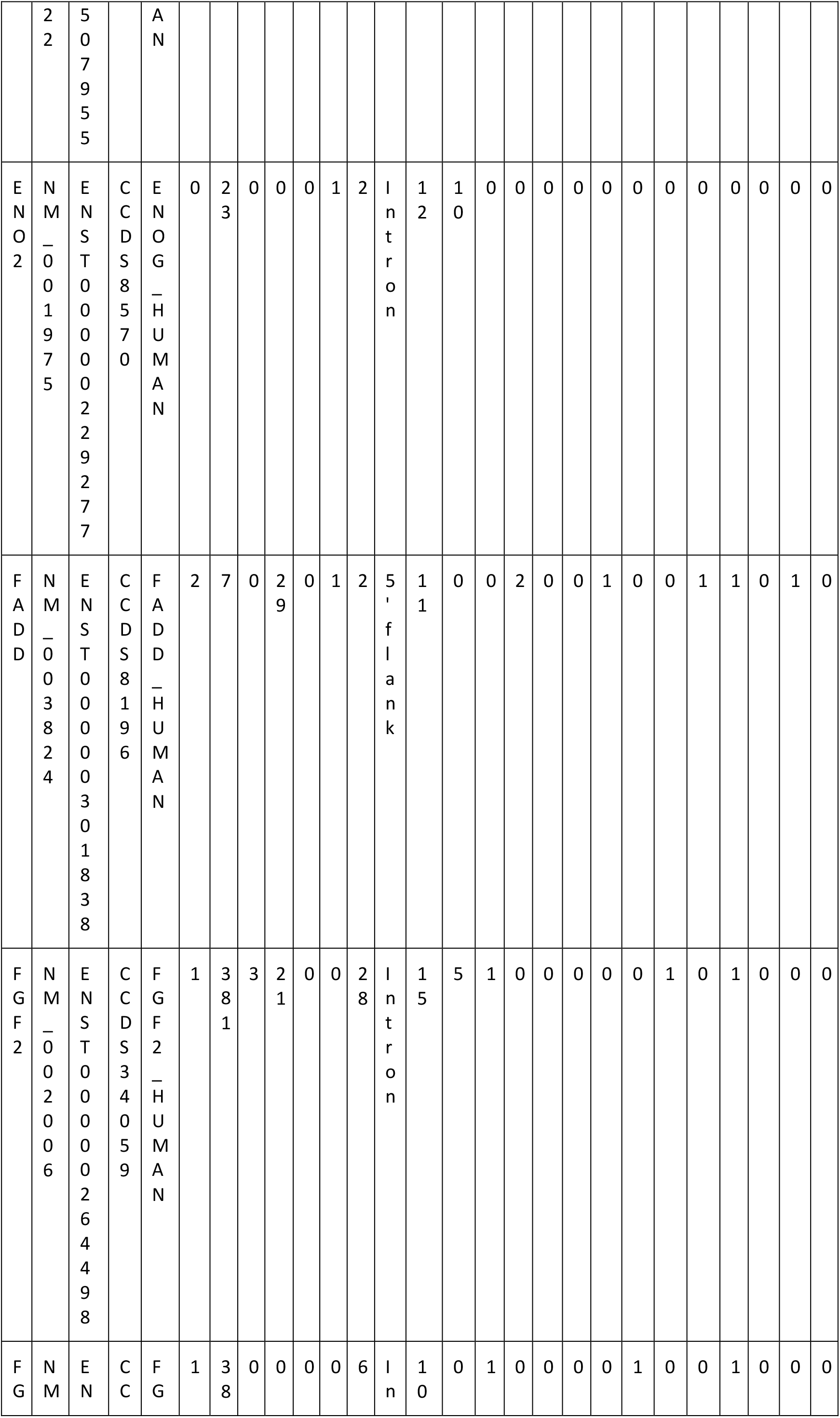
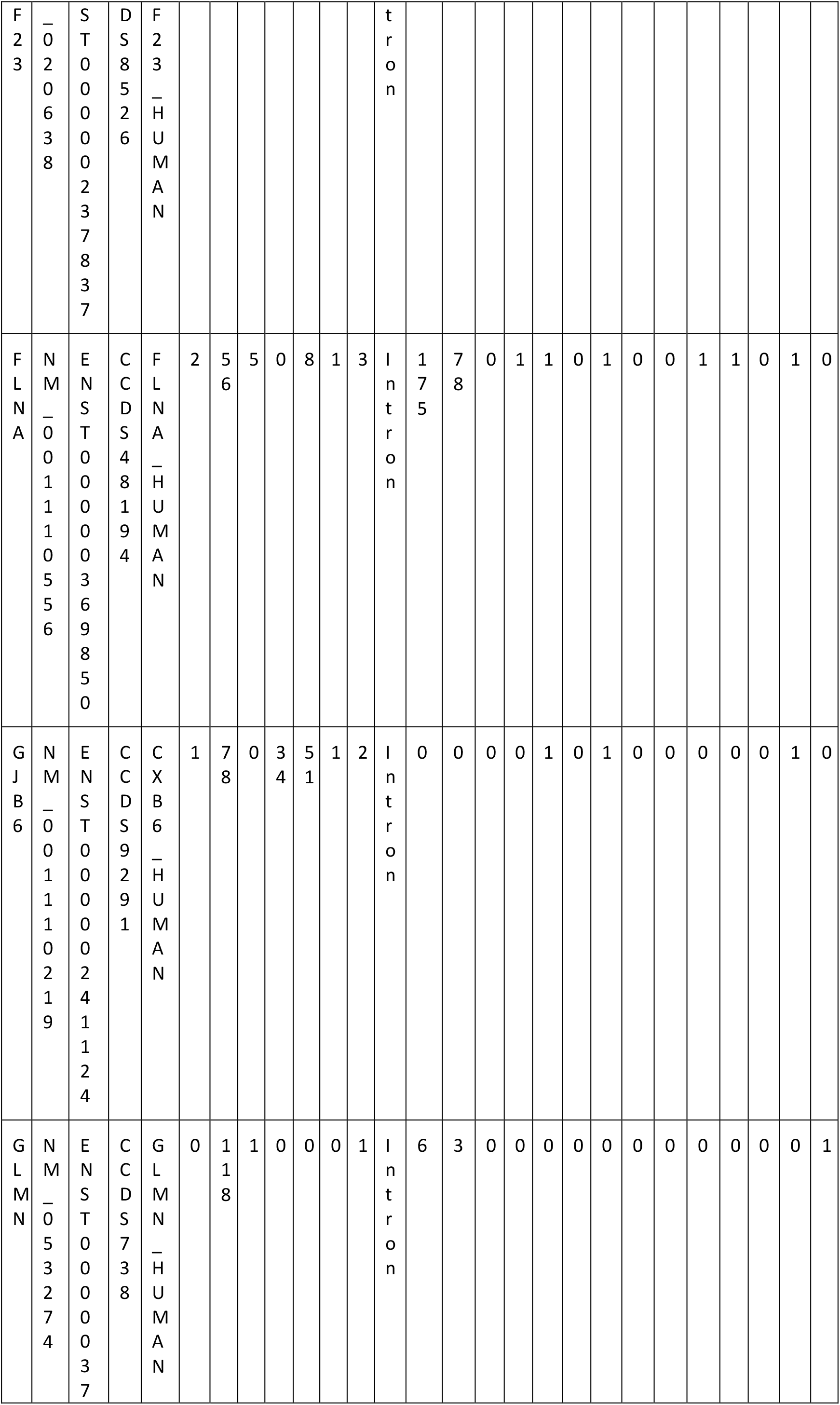
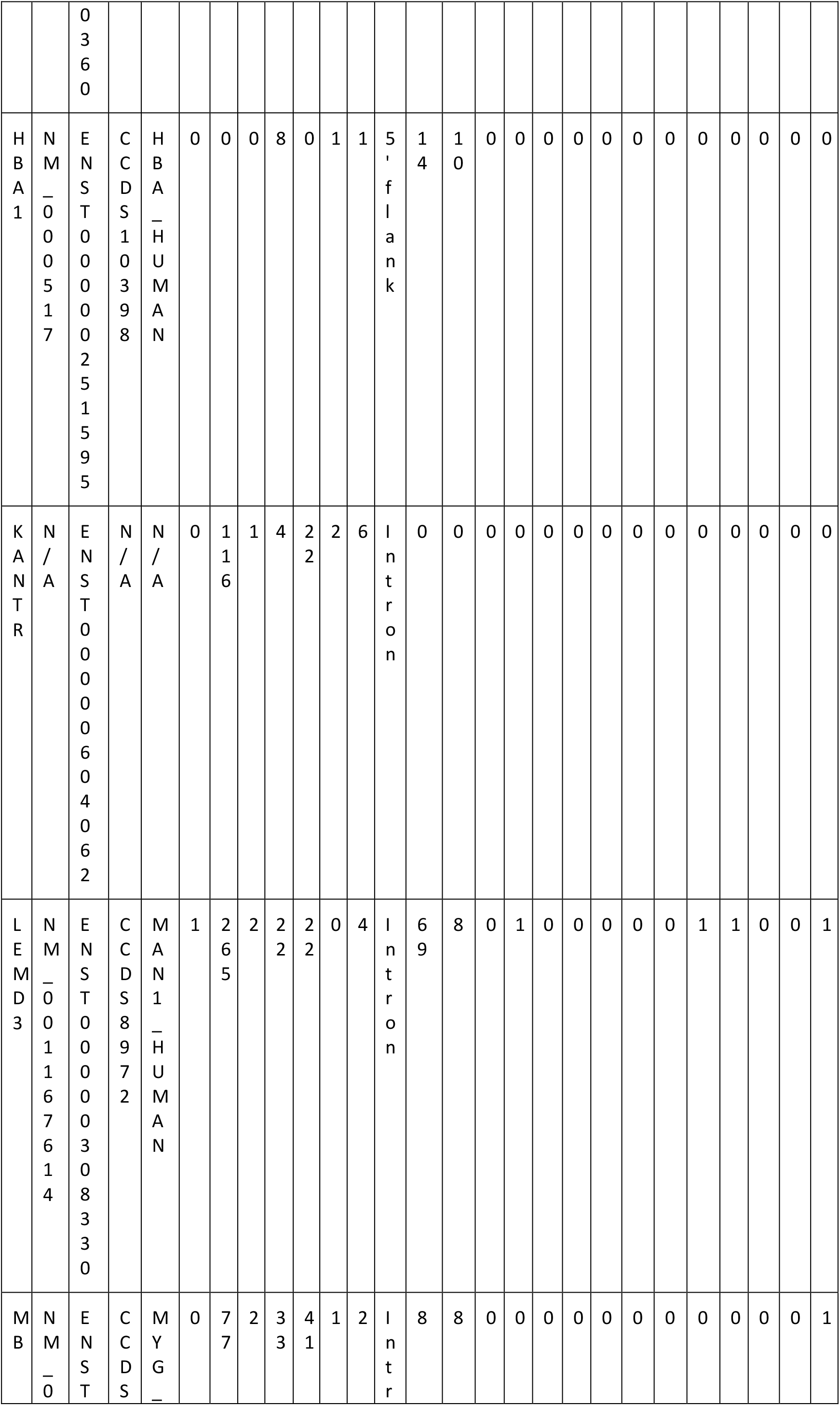
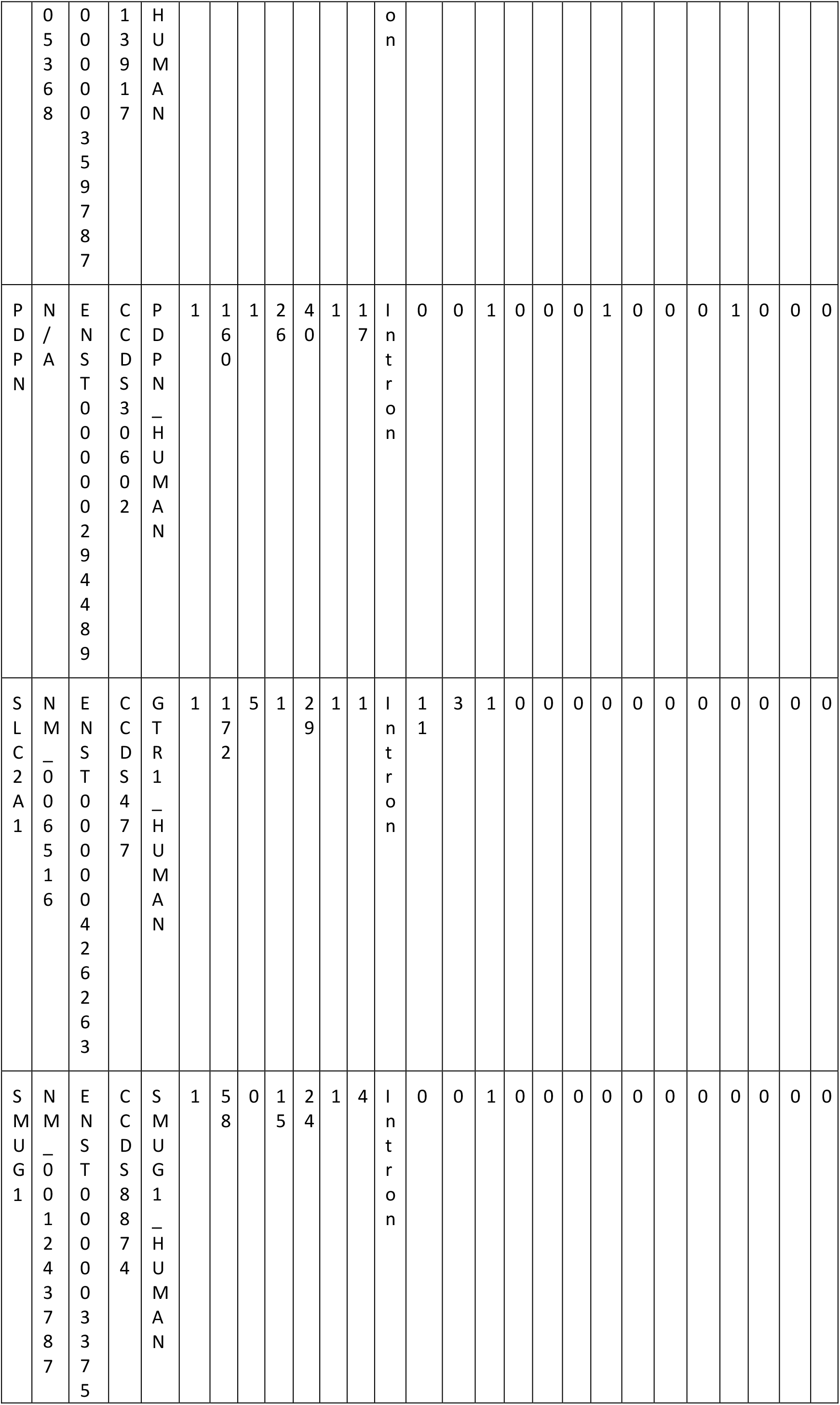
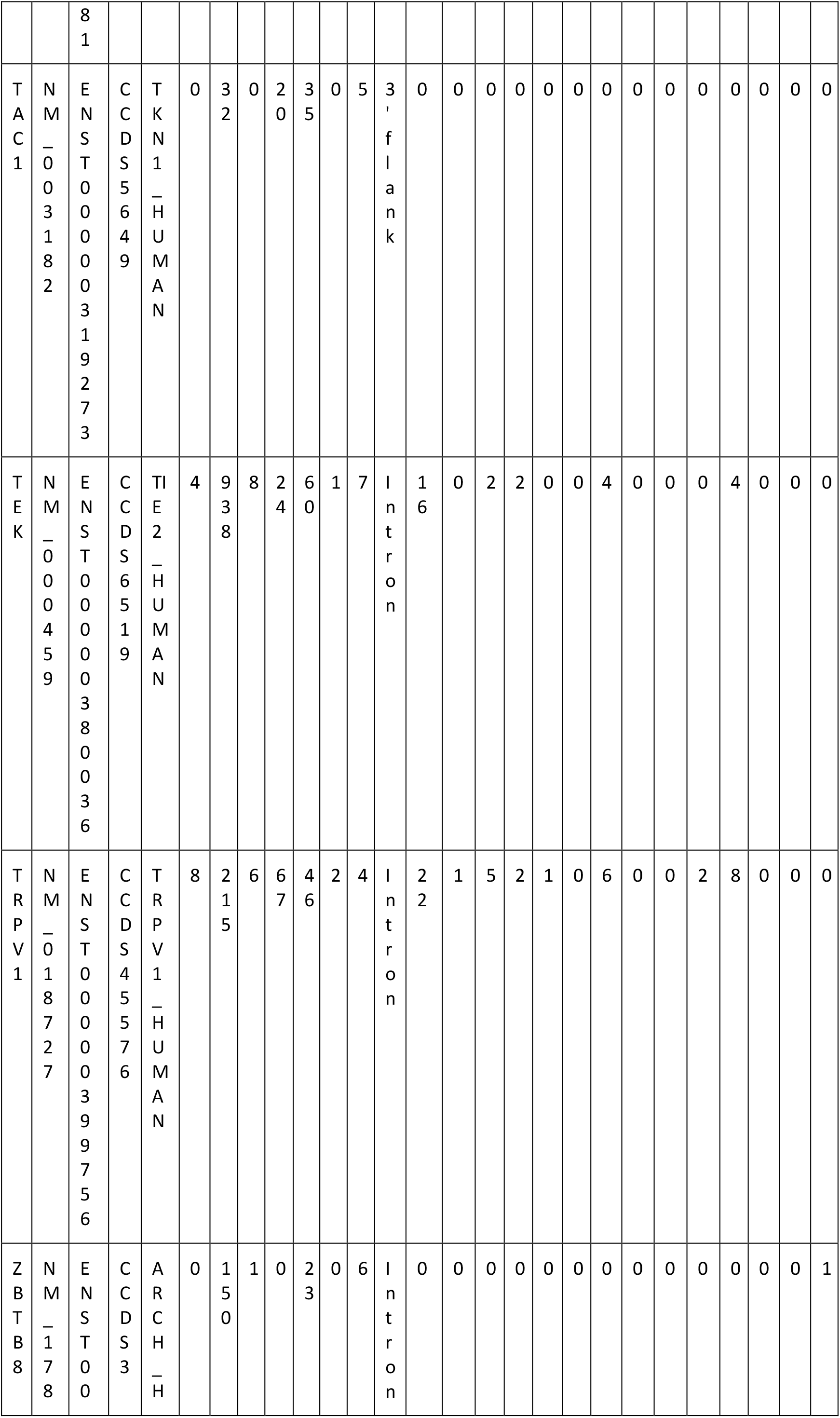
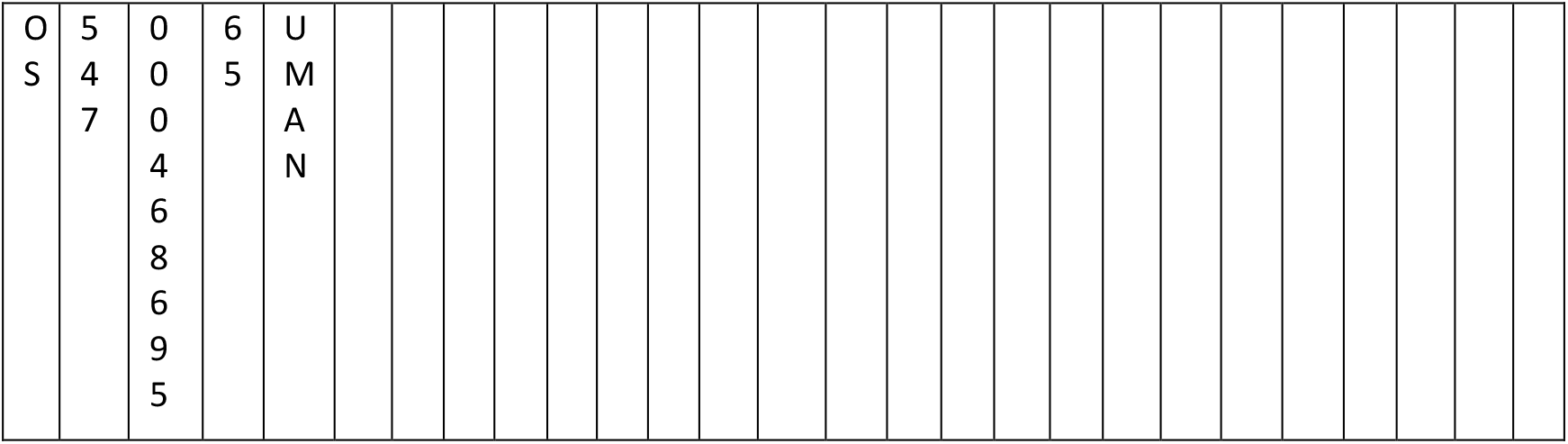
Splice mutation analysis of genes associated with other CVDs. Tabulated information includes Name, RefSeq Ensembl, CCDS, Uniprot Missense, Intron, Silent, 5’ flank, 3’ flank, 5’ UTR, 3’ UTR, Highest frequency mutation type, Phosphorylation sites, Acetylation sites, Neutral impact, Low impact, Medium impact, High impact, Tolerated, Tolerated low confidence, Deleterious low confidence, Deleterious, Benign, Possibly damaging, Probably damaging.

## 4. Discussion

To study chronic diseases with complex biological processes holistically, it is imperative to take an integrative approach that combines multi-omics data to highlight the interrelationships of the involved molecular functions. This provides information to fathom the complex biology systematically. Our approach combines clinical, gene expression and genetic variant data, in a sequential or simultaneous manner, to understand the interplay of genes in CVD. The results help in assessing the flow of information from one omics level to the other in bridging the gap from transcriptomic to genomic.

The overall goal of this study was to conduct an integrative CVD study, involving RNA-seq gene expression analysis verified through gene-variant analysis of HF and other CVD using WGS. We combined our previous findings from gene expression analysis with a gene-variant study to explore mutated genes with altered expression. Our results reinforce the results of the differential expression and variant analyses of these genes and bolster the link between the irregular expression of these genes and CVD. Further, we also noticed that mutation percentage in CVD genes was higher in HF patients compared to other CVD phenotypes (Figure 3B). The number of mutations per gene is also higher in few HF genes (Figure 3C). Additional testing of these genes may lead to the development of new clinical tools to improve diagnosis and prognosis of CVD patients.

WGS allowed us to do in-depth analysis of CVD genes as RNA-seq cannot detect any of the variants located in noncoding DNA regions. Another challenge detecting DNA variants using RNA-Seq is related to the intrinsic complexity of the computational analysis of RNA data, which requires correct mapping of RNA-seq reads to the reference human genome. Mutations observed in the altered CVD genes were mostly intronic, 5’ flank, and 3’ flank mutations. it is very difficult to prove the effect of intronic mutations except for those affecting splice sites and known regulatory sequences. If the identified intronic alterations are in regulatory regions, they could affect gene expression levels. Point mutations in introns can also introduce novel splice sites causing transcriptional and translational regulation, activate novel promoters, or introduce/eliminate enhancer activity. Insertions/deletions (indels) in intronic areas may carry even more dramatic effects.

To perform and support translational research, it is important to connect genomic findings with the medical records of the patients who consented and participated in this study. However, the challenge is to integrate clinical and genomic data of variable lengths and volumes to be useful in clinical settings. To do this, we needed to reformat information such that it could be integrated and linked to electronic health systems (e.g., EPIC, NextGen, Cerner, etc.). In health systems all around the world, diseases/diagnosis are represented as the International Classification of Diseases (ICD) codes. These codes are maintained by the World Health Organization (WHO). Unlike ICD codes, genomics information has not been internationally standardized; however, some data are very well maintained, such as in Ensembl [37] and Gencode [38]. In this study, we linked ICD-10 codes with CVD genes. Among genes associated with HF, 29 were associated with congestive heart failure (code I50.9), 8 were associated with diastolic heart failure (code I50.3), and 11 were associated with systolic heart failure (code I50.20). For genes associated with other CVDs, 19 were associated with the ICD-10 cardiovascular organ benign neoplasm (code D15.1), 3 were associated with cardiovascular syphilis (A52.00), and 1 was associated with Infections recurrent with encephalopathy hepatic dysfunction and cardiovascular malformations (D53.0). Details are provided in Table 3.

**Table 3.** Linked ICD-10 codes with all the genes investigated for being associated with HF and other CVDs. Information includes Gene name, Ensembl ID, Genecode, ICD-10 Code, and Disease.

| Gene | Ensembl Id | Genecode | ICD-10 | Disease |
| --- | --- | --- | --- | --- |
| TNF | ENSG00000232810 | ENSG00000232810.3 | I50.20 | Systolic heart failure |
| IL6 | ENSG00000136244 | ENSG00000136244.11 | I50.20 | Systolic heart failure |
| ACE | ENSG00000159640 | ENSG00000159640.15 | I50.9 | Congestive heart failure |
| ACE | ENSG00000159640 | ENSG00000159640.15 | I50.3 | Diastolic heart failure |
| ACE | ENSG00000159640 | ENSG00000159640.15 | I50.20 | Systolic heart failure |
| MMP2 | ENSG00000087245 | ENSG00000087245.12 | I50.3 | Diastolic heart failure |
| NOS3 | ENSG00000164867 | ENSG00000164867.10 | I50.3 | Diastolic heart failure |
| AGT | ENSG00000135744 | ENSG00000135744.7 | I50.3 | Diastolic heart failure |
| EDN1 | ENSG00000078401 | ENSG00000078401.6 | I50.9 | Congestive heart failure |
| REN | ENSG00000143839 | ENSG00000143839.14 | I50.9 | Congestive heart failure |
| MYH7 | ENSG00000092054 | ENSG00000092054.12 | I50.9 | Congestive heart failure |
| AGTR1 | ENSG00000144891 | ENSG00000144891.17 | I50.3 | Diastolic heart failure |
| AGTR1 | ENSG00000144891 | ENSG00000144891.17 | I50.9 | Congestive heart failure |
| NPPA | ENSG00000175206 | ENSG00000175206.10 | I50.9 | Congestive heart failure |
| ADRB2 | ENSG00000169252 | ENSG00000169252.5 | I50.9 | Congestive heart failure |
| NR3C2 | ENSG00000151623 | ENSG00000151623.14 | I50.9 | Congestive heart failure |
| NR3C2 | ENSG00000151623 | ENSG00000151623.14 | I50.20 | Systolic heart failure |
| MME | ENSG00000196549 | ENSG00000196549.10 | I50.9 | Congestive heart failure |
| CRP | ENSG00000132693 | ENSG00000132693.12 | I50.20 | Systolic heart failure |
| MYH6 | ENSG00000197616 | ENSG00000197616.11 | I50.9 | Congestive heart failure |
| EPO | ENSG00000130427 | ENSG00000130427.2 | I50.9 | Congestive heart failure |
| CST3 | ENSG00000101439 | ENSG00000101439.8 | I50.20 | Systolic heart failure |
| EDNRA | ENSG00000151617 | ENSG00000151617.15 | I50.9 | Congestive heart failure |
| AQP2 | ENSG00000167580 | ENSG00000167580.7 | I50.9 | Congestive heart failure |
| MYBPC3 | ENSG00000134571 | ENSG00000134571.10 | I50.3 | Diastolic heart failure |
| KNG1 | ENSG00000113889 | ENSG00000113889.12 | I50.9 | Congestive heart failure |
| VCL | ENSG00000035403 | ENSG00000035403.17 | I50.9 | Congestive heart failure |
| HOTAIR | ENSG00000228630 | ENSG00000228630.5 | I50.9 | Congestive heart failure |
| CDKN2B-AS1 | ENSG00000240498 | ENSG00000240498.7 | I50.9 | Congestive heart failure |
| ANKRD1 | ENSG00000148677 | ENSG00000148677.6 | I50.3 | Diastolic heart failure |
| ADM | ENSG00000148926 | ENSG00000148926.9 | I50.9 | Congestive heart failure |
| AMPD1 | ENSG00000116748 | ENSG00000116748.21 | I50.9 | Congestive heart failure |
| PLN | ENSG00000198523 | ENSG00000198523.5 | I50.9 | Congestive heart failure |
| LGALS3 | ENSG00000131981 | ENSG00000131981.15 | I50.20 | Systolic heart failure |
| NPPB | ENSG00000120937 | ENSG00000120937.8 | I50.9 | Congestive heart failure |
| NPPB | ENSG00000120937 | ENSG00000120937.8 | I50.3 | Diastolic heart failure |
| NPPB | ENSG00000120937 | ENSG00000120937.8 | I50.20 | Systolic heart failure |
| ADRB1 | ENSG00000043591 | ENSG00000043591.5 | I50.9 | Congestive heart failure |
| ADRB1 | ENSG00000043591 | ENSG00000043591.5 | I50.20 | Systolic heart failure |
| UTS2 | ENSG00000049247 | ENSG00000049247.13 | I50.9 | Congestive heart failure |
| PIK3C2A | ENSG00000011405 | ENSG00000011405.13 | I50.9 | Congestive heart failure |
| NPPC | ENSG00000163273 | ENSG00000163273.3 | I50.9 | Congestive heart failure |
| CORIN | ENSG00000145244 | ENSG00000145244.11 | I50.20 | Systolic heart failure |
| NPR1 | ENSG00000169418 | ENSG00000169418.9 | I50.9 | Congestive heart failure |
| LSINCT5 | ENSG00000281560 | ENSG00000281560.1 | I50.9 | Congestive heart failure |
| TUSC7 | ENSG00000243197 | NA | I50.9 | Congestive heart failure |
| HSPB7 | ENSG00000173641 | ENSG00000173641.17 | I50.20 | Systolic heart failure |
| RP11-451G4.2 | ENSG00000240045 | ENSG00000240045.1 | I50.9 | Heart failure |
| SLC2A1 | ENSG00000117394 | ENSG00000117394.21 | D15.1 | Cardiovascular organ benign neoplasm |
| FGF2 | ENSG00000138685 | ENSG00000138685.14 | D15.1 | Cardiovascular organ benign neoplasm |
| FLNA | ENSG00000196924 | ENSG00000196924.15 | D15.1 | Cardiovascular organ benign neoplasm |
| HBA1 | ENSG00000206172 | ENSG00000206172.8 | D15.1 | Cardiovascular organ benign neoplasm |
| GJB6 | ENSG00000121742 | ENSG00000121742.17 | D15.1 | Cardiovascular organ benign neoplasm |
| ATP2A2 | ENSG00000174437 | ENSG00000174437.16 | D15.1 | Cardiovascular organ benign neoplasm |
| CD40LG | ENSG00000102245 | ENSG00000102245.7 | A52.00 | Cardiovascular syphilis |
| FGF23 | ENSG00000118972 | ENSG00000118972.1 | D15.1 | Cardiovascular organ benign neoplasm |
| TEK | ENSG00000120156 | ENSG00000120156.20 | D15.1 | Cardiovascular organ benign neoplasm |
| TAC1 | ENSG00000006128 | ENSG00000006128.11 | D15.1 | Cardiovascular organ benign neoplasm |
| DDX41 | ENSG00000183258 | ENSG00000183258.11 | A52.00 | Cardiovascular syphilis |
| FADD | ENSG00000168040 | ENSG00000168040.4 | D53.0 | Infections recurrent with encephalopathy hepatic dysfunction and cardiovascular malformations |
| ENO2 | ENSG00000111674 | ENSG00000111674.8 | D15.1 | Cardiovascular organ benign neoplasm |
| LEMD3 | ENSG00000174106 | ENSG00000174106.2 | D15.1 | Cardiovascular organ benign neoplasm |
| CD34 | ENSG00000174059 | ENSG00000174059.16 | D15.1 | Cardiovascular organ benign neoplasm |
| TRPV1 | ENSG00000196689 | ENSG00000196689.11 | D15.1 | Cardiovascular organ benign neoplasm |
| GLMN | ENSG00000174842 | ENSG00000174842.16 | D15.1 | Cardiovascular organ benign neoplasm |
| MB | ENSG00000198125 | ENSG00000198125.12 | D15.1 | Cardiovascular organ benign neoplasm |
| SMUG1 | ENSG00000123415 | ENSG00000123415.15 | A52.00 | Cardiovascular syphilis |
| PDPN | ENSG00000162493 | ENSG00000162493.16 | D15.1 | Cardiovascular organ benign neoplasm |
| CALD1 | ENSG00000122786 | ENSG00000122786.19 | D15.1 | Cardiovascular organ benign neoplasm |
| KANTR | ENSG00000232593 | NA | D15.1 | Cardiovascular organ benign neoplasm |
| ZBTB8OS | ENSG00000176261 | ENSG00000176261.15 | D15.1 | Cardiovascular organ benign neoplasm |

Integrative approaches, by virtue of their ability to study biological phenomena holistically, could improve prognostics and predictive accuracy of disease phenotypes, and hence can eventually aid in better treatment and prevention. In the future, we seek to evaluate the causal basis for HF and other CVDs by moving beyond the one gene-one disease model through the integration of the expressed genome, and characterization of co-morbidities and phenotypic traits derived from metabolomic signatures and EHR. We aim to contribute to the paradigm shift in the application and interpretation of genetic and genomically informed medicine for HF and other CVDs, moving from a deterministic conceptualization to a probabilistic interpretation of genetic risk. This will support diagnostic and preventive care delivery strategies beyond traditional symptom-driven, disease-causal medical practice. By complementing RNAseq with mutation load, we aim to identify unifying and indication-specific molecular profiles that can predict response to treatment. Machine learning models can help us identify a baseline transcriptional signature highly predictive of response across these indications. This will accelerate our ability to leverage and extend the information contained within the original data, and to model patient-specific genomics and clinical data for significant transcriptional correlate of response highlighting the association of genetic variants to outcome of treatment in HF and other CVD mechanisms [17].

We are also interested in elucidating how changes in gene expression regulation through alternative splicing contribute to CVD. Alternative splicing is a key control point in gene expression, and its misregulation often leads to malignancy. However, most studies on splicing have not fully profiled splicing in CVD [39]. In future, we aim to explicate isoform-level diversity in CVD with a subset of the novel spliced isoforms with biological significance that may serve as potential biomarkers for innovative therapeutics. Although vast progress has been made in understanding the global “omics” architecture of CVD, there remains a considerable gap in understanding population differences in minorities and under-represented populations. However, to date, no one has examined differences in CVD-specific splicing events between racial groups, and their impact on health disparities, or the association between CVD specific splicing events and certain external exposures, such as smoking.

## Acknowledgements

We appreciate great support by the Pat and Jim Calhoun Cardiology Center, and Department of Genetics and Genome Sciences, at the UConn School of Medicine, UConn Health; Rutgers Institute for Health, Health Care Policy and Aging Research (IFH), and Rutgers Robert Wood Johnson Medical School (RWJMS), Rutgers Biomedical and Health Sciences (RBHS) at the Rutgers, The State University of New Jersey.

We would like to give special thanks to Dr. Christopher Bonin and Dr. Geneva Hargis for stylistic and native speaker corrections.

## Author contributions

ZA lead this study. ZA did RNA-seq data processing, quality checking, and downstream analysis. ZA developed MAV-clic and PROMIS-LCR, supervised JWES and GVViZ implementation, and performed cohort building and integrative clinical data analysis of consented patients. SZ provided bioinformatics expertise in variant data analysis and visualization. NP supported post-computational analysis and evaluation of results. ZA drafted the paper. All authors have participated in writing and review, and have approved paper for publication. BL proposed, supervised, and supported the study.

## Authors’ information

ZA is the Assistant Professor of Medicine – Tenure Track and Core Member at the Rutgers Institute for Health, Health Care Policy and Aging Research; and Department of Medicine – Division of General Internal Medicine, Rutgers Robert Wood Johnson Medical School, Rutgers Biomedical and Health Sciences, Rutgers University-New Brunswick. ZA is the Adjunct Assistant Professor at the Department of Genetics and Genome Sciences, UConn School of Medicine, UConn Health, CT; and Full Academic Member of the Rutgers Microbiology and Molecular Genetics; Center for Cancer Health Equity, Rutgers Cancer Institute of New Jersey; Rutgers Human Genetics Institute of New Jersey, NJ.

NP is the MD student at the Rutgers Robert Wood Johnson Medical School.

SZ is the Senior Scientist Associated with the Rutgers Cancer Institute of New Jersey.

BL is the Interim Chief Executive Officer, UConn Health; Executive Vice President for Health Affairs; Dean, UConn School of Medicine; Director, Pat and Jim Calhoun Cardiology Center; and Ray Neag Distinguished Professor of Cardiovascular Biology and Medicine. BL is an internationally recognized cardiovascular physician-scientist and national leader in academic medicine.

## Declarations

### Ethical Approval and Consent to participate

Informed consent was obtained from all subjects. All human samples were used in accordance with relevant guidelines and regulations, and all experimental protocols were approved by Institutional Review Board, UConn Health.

### Consent for publication

Not applicable

### Availability of data and material

Attached as tables.

### Competing interests

The Authors declare no Competing Financial or Non-Financial Interests.

### Funding

The study was supported by the School of Medicine, UConn Health, CT.

### Data Availability Statement

The data analyzed in the current study are attached, and available from the corresponding author on reasonable request.

## Supplementary information

Accompanies this paper:

### Supplementary material 1

Heart Failure and Cardiovascular Diseases (CVD) Gene-Variant Analysis.

## Notes

### Competing Interest Statement

The authors have declared no competing interest.

