## Supplementary Material for "Investigating variant and expression of CVD genes associated phenotypes among high-risk Heart Failure patients"

Heart Failure and Cardiovascular Diseases (CVD) Gene-Variant Analysis

### 1. Mutation analysis of HF genes

**ACE:** An enzyme that converts angiotensin I into angiotensin II and inactivates bradykinin, both of which ultimately cause an increase in blood pressure [1]. Certain mutations (silent and benign or harmful) in the ACE gene have been shown to increase the levels of ACE in individuals and lead to CVDs such as a myocardial infarction [2]. Specifically, a deletion allele also called the D allele has been associated with HF, high blood pressure, and non-CVDs such as complications with diabetes and asthma [3]. We generated a lollipop plot that mapped mutations identified in the literature onto specific positions of the amino acid sequence of ACE (Figure 4A). A total of 1033 mutations were mapped onto the sequence, 6 of which are missense mutations, 23 of which are splice mutations, 757 which are intron mutations, 115 which are 5' flank mutations, 42 which are 3' UTR mutations, and 90 which are silent. We found a total of 6 phosphorylation sites, 2 acetylation sites, 1 ubiquitination site, and 13 N-linked glycosylation sites. Six missense mutations were found to have some negative impact on the function of the protein: a couple of missense mutations were mapped at Y244C, and other missense mutations were mapped to A261S, P485R, S660C. These all had medium functional impact, scored by mutation assessor, but only one mapped to D592G with low functional impact (score of deleterious impact by SIFT, and a score of possibly damaging by PolyPhen-2). We found 25 splice mutations, one mapped to X529 and the other to X1127, and all remaining (23) splice mutations mapped to X1231.

**ADM:** A hormone produced by multiple tissues including the adrenal medulla and the heart. ADM plays a role in controlling blood pressure because it widens blood vessels which causes blood pressure to decline [4]. We plotted known mutations of ADM onto the amino acid sequence and found 359 mapped mutations (Figure 4B). A couple were intron mutations, 189 were 5' flank mutations, 143 were 3' flank mutations, 1 was a 5' UTR mutation, and 24 were 3' UTR mutations. No information regarding the functional impact of any of these mutations was provided. There was a total of 9 phosphorylation sites, 2 amidation sites, and 11 O-linked glycosylation sites. For ADM, 3' and 5' flank mutations were the most numerous mutations, and these both 5' and 3' flank regions have been known to increase transcription [90] and this may relate to the fact that ADM is found to be high in a few patients with HF and CVDs.

**ADRB1:** A receptor found primarily in the heart. Compounds such as epinephrine and norepinephrine bind to ADRB1 and can cause both heart rate and contractility to increase [5]. We found a total of 431 mutations in ARDB1 (Figure 4C), which were mapped onto the amino acid sequence. 44 were missense mutations, 1 was a silent mutation, 179 were 5'f lank mutations, 181 3' flank mutations, 2 were 5' UTR mutations, and 24 were 3' UTR mutations. Of these 431 mutations, only one was found to be deleterious, which was found at position C393G. The rest of the mutations were classified as neutral impact, low impact, tolerated, or benign. There was no intron mutation found in ADRB1, and a total of 12 phosphorylation sites and no other post-translational modifications (PTM). 5' and 3' flank mutations were found to be the most numerous mutations mapped to ADRB1.

**ADRB2:** A receptor whose function is like ADRB1, but it also affects vascular smooth muscle and bronchial smooth muscle by causing both to widen [5]. We found and plotted a total of 837 mutations in ADRB2 (Figure 4D). All the functional impact scores of the mutations are low impact. 45 mutations were missense mutations, 72 were silent, 575 were 5' flank mutations, 67 were 3' flank mutations, 52 were 5' UTR mutations, and 26 were 3' UTR mutations. Like ADRB1, ADRB2 had no intron mutations mapped onto its sequence. There was a total of 24 phosphorylation sites, 2 hydroxylation sites, and 1 palmitoylation site.

**AGT:** A protein that plays a major role in the renin-angiotensin system because it is used to ultimately produce angiotensin II through various enzymes [6]. Angiotensin II increases blood pressure and AGT is necessary for this effect to occur [7, 8]. We found and plotted a total of 1070 mutations onto the AGT amino acid sequence (Figure 4E). 33 of the mutations were missense, 502 were intron mutations, 34 were silent mutations, 315 were 5' flank mutations, 163 were 3' flank mutations, and 23 were 3' UTR mutations. Most of the mutations were mapped onto AGT were intron. 1070 mutations were mapped onto AGT, including 24 with neutral impact, 9 with medium impact, 24 tolerated, 24 benign, 9 are deleterious, and 9 damaging mutations. There are a total of 3 phosphorylation sites, 1 acetylation site, and 6 N-linked glycosylation sites.

**AGTR1:** A protein that serves as the target receptor for angiotensin II and causes an increase in blood pressure [9]. Elevated activity of AGTR1 has been associated with many cardiovascular diseases including high blood pressure and atherosclerosis [10]. We found and plotted a total of 2487 mutations on the AGTR1 amino acid sequence (Figure 4F). None of these mutations were missense. A total of 1785 are intron mutations, followed by 3' and 5' flank mutations. 32 are silent mutations, 245 are 5' flank mutations, 337 are 3' flank mutations, and 23 are 3’ UTR mutations. In terms of PTM’s, there are a total of 10 phosphorylation sites, 1 N-linked glycosylation site, and 1-S nitrosylation site.

**AMPD1:** A gene that codes for a protein that is imperative to the normal function of skeletal muscle and a specific mutation in AMPD1 has been known to lead to musculoskeletal problems [11]. Interestingly, this same mutation has also been shown to have beneficial cardiovascular effects such as alleviating symptoms associated with HF. A total of 800 mutations were found in AMPD1 (Figure 4G) including: 8 are missense mutations, 4 are truncating/nonsense mutations, 661 are intron mutations, 1 is a silent mutation, and 126 are 5’ flank mutations. Amongst these mutations, 1 has low impact, 4 have medium impact, 3 have high impact, 1 is tolerated with low confidence, 7 are deleterious, and 8 are damaging. In terms of PTM’s, there were 9 phosphorylation sites and 2 acetylation sites. Most of the mutations were mapped in intron, followed by 5’ flank mutations.

**ANKRD1:** A gene that codes for a protein called cardiac ankyrin repeating protein (CARP). Increased levels of CARP have been linked to HF [12]. We found a total of 428 mutations onto the sequence of ANKDR1 (Figure 4H): 148 are intron mutations, 105 are 5’ flank mutations, 144 are 3’ flank mutations, and 31 are 3’ UTR mutations. No information about the functional impact of these mutations was available. In terms of PTM’s, there were only 3 phosphorylation sites.

**AQP2:** A protein channel that plays a role in storing water in the body via the kidneys and is heavily influenced by anti-diuretic hormone [13, 14]. During variant analysis, we found 793 mutations onto the AQP2 amino acid sequence (Figure 4I). There are a total of 10 splice mutations, 423 intron mutations, 34 silent mutations, 142 5’ flank mutations, and 184 3’ UTR mutations. Most of mutations are in intron, followed by 3’ UTR and 5’ flank mutations. In terms of PTM’s, there are a total of 6 phosphorylation sites and 1 ubiquinitation site.

**CORIN:** A protein which functions to generate atrial natriuretic peptide (ANP) hormone to lower blood pressure, and low levels of the ANP is linked with high blood pressure [15, 16]. A total of 9566 mutations were mapped onto the CORIN sequence (Figure 4J). Of these mutations, 71 are missense, 9413 are intron mutations, 32 are silent, 2 are 3’ flank, and 42 are 3’ UTR mutations. Among all, 69 were classified as having a neutral functional impact, 1 is classified as having a low impact, 1 was classified as having a medium impact, 36 are tolerated, 33 are tolerated low confidence, 1 is deleterious low confidence, 1 is deleterious, 70 are benign, and 1 is possibly damaging. Majority of the mutations that do have a functional impact score tend to be on the lower end of the spectrum. In terms of PTM’s, there are only 3 phosphorylation sites. Most of the mutations are in intron sequence and have relatively few 5' and 3' flank mutations.

**CRP:** A protein that is made by liver cells, and increased level of CRP has been associated with CVDs [17]. We found and plotted 403 mutations in CRP (Figure 4K): 1 is a missense mutation, 24 are intron mutations, 4 are silent mutations, 277 are 5' flank mutations, 60 are 3' flank mutations, and 37 are 3' UTR mutations. In terms of PTM's, there was 1 methylation site and 1 pyrrolidone carboxylic acid site. Of these mutations, only the 1 missense mutation had a functional impact score from each of the 3 programs: a medium functional impact, a deleterious impact, and a possibly damaging impact. Most of the mutations are 5' flank followed by the 3' flank.

**CST3:** A protein that controls the levels of cathepsin proteins. Its high levels have been associated with many CVDs including heart failure and stroke [18, 19, 20]. We mapped a total of 1386 mutations on the CST3 sequence (Figure 4L): 11 are missense, 123 are intron, 38 are silent, 387 are 5' flank, 687 are 3' flank, 13 are 5' UTR, and 127 are 3' UTR mutations. In terms of PTM's, there is only one phosphorylation site. The most numerous mutations mapped onto the sequence are 3' flank mutations followed by 5' flank mutations.

**EDN1:** A protein that causes blood vessels to narrow and widen based on the receptor it binds [21]. A total of 871 mutations were found in EDN1 (Figure 4M): 17 are missense, 359 are intron, 35 are silent, 269 are 5' flank, 187 are 3' flank, and 4 are 3' UTR mutations. In terms of PTM's, there are 3 phosphorylation sites, including 1 O-linked glycosylation site, and 2 methylation sites. Most of the numerous mutations mapped onto the EDN1 sequence are intron mutations followed by 5' and 3' flank.

**EDNRA:** A protein receptor found primarily in the vasculature which elicits narrowing of blood vessels whenever its agonist, endothelin-1, binds to it [22]. Its increased levels are linked to HF. We found and mapped a total of 1833 mutations onto the EDNRA sequence (Figure 4N): 1 of these mutations is a splice mutation, 1476 mutations are intron mutations, 32 are silent, 113 are 5' flank mutations, 113 are 3' flank mutations, 19 are 5' UTR mutations, and 79 are 3' UTR mutations. In terms of PTM's, there are 18 phosphorylation sites and 1 acetylation site. Most of the mutations were mapped to introns, followed by the 5' and 3' flank mutations.

**EPO:** A hormone that promotes the synthesis of red blood cells. Its abnormal amounts in patients have been linked to HF [23, 24]. We found and mapped a total of 537 mutations onto the sequence of EPO (Figure 4O): 1 of these mutations is a splice mutation, 50 are intron mutations, 361 are 5’ flank mutations, 84 are 3’ flank mutation, and 41 are 3’ UTR mutations. In terms of PTM’s, there are 4 phosphorylation sites, 6 N-linked glycosylation sites, and 1 O-linked glycosylation site. Most of the mutations are mapped to 5’ flank mutations followed by 3’ flank and intron mutations.

**HSPB7:** A protein that affects actin formation and is found in the heart, and mutations in this gene have been linked to HF [25]. We found and mapped a total of 1234 mutations on the protein sequence (Figure 4P): 364 are intron mutations, 87 are silent mutations, 231 are 5’ flank mutations, 313 are 3’ flank mutations, 37 are 5’ UTR mutations, and 202 are 3’ UTR mutations. In terms of PTM’s, there are only 6 phosphorylation sites. Most of the numerous mutations are in intron, followed by 3’ and 5’ flank mutations.

**IL6:** A cytokine protein that stimulates inflammation. High levels of IL6 have been associated with HF [26, 27]. We found and mapped a total of 388 mutations onto the IL6 sequence (Figure 4Q): 227 are intron mutations, 1 is a 5’ flank mutation, 159 are 3’ flank, and 1 is a 3’ UTR mutation. In terms of PTM’s, there are a total of 3 phosphorylation sites, 2 N-linked glycosylation sites, and 3 are O-linked glycosylation sites. Most mutations mapped onto the sequence are intron mutations followed by 3’ flank mutations whose role in IL6 functioning has not have been defined yet.

**KNG1:** A gene that generates proteins that effect blood pressure and mutations in KNG1 are linked to high blood pressure and HF [28]. We found and mapped a total of 208 mutations onto the sequence (Figure 4R): 31 are missense mutations, 3 are truncating mutations, 111 are intron mutations, 43 are silent mutations, 19 are 3’ flank mutations, and 1 is a 3’ UTR mutation. Of these mutations, 46 have a neutral impact, 6 have a medium impact, 25 are tolerated, 6 are deleterious, 25 are benign, and 6 are possibly damaging. In terms of PTM’s, there are 11 phosphorylation sites, 2 acetylation sites, 4 N-linked glycosylation sites, 1 O-linked glycosylation site, and 1 hydroxylation site. Most of the mutations mapped onto the sequence are intron mutations.

**LGALS3:** A protein released by macrophages and its high levels have been associated with HF and heart attack [29]. We discovered and mapped a total of 1154 mutations onto the sequence of LGALS3 (Figure 4S): 58 of the mutations are missense, 592 are intron, 1 is a silent, 502 are 5’ flank, and 1 is a 5’ UTR mutation. Of these mutations, 27 have a neutral impact, 31 have a medium impact, 27 are tolerated, 26 are deleterious, 27 are benign, and 26 are probably damaging. In terms of PTM’s, there are a total of 10 phosphorylation sites, 2 acetylation sites, and 3 malonylation sites. Most of the mutations mapped onto the sequence are intron mutations followed by 5’ flank mutations.

**MME:** A protein that decreases levels of ANP and its high levels have proven to be associated with heart disease [30]. We found and plotted a total of 4191 mutations onto the MME sequence (Figure 4T): 3 of the mutations are missense, 4 are splice mutations, 3820 are intron mutations, 3 are silent mutations, 238 are 5’ flank mutations, 2 are 5’ UTR mutations, and 121 are 3’ UTR mutations. Of these mutations 1 has a neutral impact, 2 have a low impact, 1 is tolerated, 2 are deleterious, and 3 are benign. In terms of PTM’s, there are 10 phosphorylation sites, 2 are acetylation sites, 4 are N-linked glycosylation sites, and 1 is a myristoylation site. Most of the mutations mapped onto the sequence are intron mutations, followed by 5’ flank mutations.

**MMP2:** A protein that breaks down parts of the extracellular matrix, and its high amounts have been shown to be associated with HF [31]. A total of 4900 mutations were mapped onto the MMP2 sequence (Figure 4U). 1 is a missense mutation, 51 are splice mutations, 4177 are intron mutations, 113 are silent, 450 are 5’ flank mutations, 81 are 5’ UTR mutations, and 27 are 3’ UTR mutations. Of these mutations, only the 1 missense mutation received functional impact scores. It was rated as having low and tolerated scores but was rated as probably damaging by the Polyphen 2 program. In terms of PTM’s, there are a total of 14 phosphorylation sites. Majority of these mutations mapped onto the sequence are intron mutations followed by 5’ flank mutations.

**MYBPC3:** A protein found in heart muscle to contract, and with a mutation, is proven to be linked HF [32]. A total of 872 mutations were mapped onto the MYBPC3 sequence (Figure 4V): 23 are missense mutations, 646 are intron mutations, 34 are silent, 107 are 5’ flank mutations, and 62 are 3’ flank mutations. Of these mutations, 8 have neutral impact, 1 has low impact, 12 have a medium impact, 2 have a high impact, 8 are tolerated, 4 are deleterious, 9 are benign, and 3 are probably damaging. In terms of PTM’s, there are 34 phosphorylation sites and 1 acetylation site. Most of the common mutations were mapped to intron, followed by the 5’ flank mutation.

**MYH6:** Found as a component of the myosin protein in the heart, and having a mutation is associated with HF, and dysfunction [33]. We found and plotted a total of 764 mutations mapped onto the MYH6 sequence (Figure 4W): 26 are missense mutations, 608 are intron mutations, 75 are silent, and 55 are 5’ flank mutations. Of these mutations 19 are neutral mutations, 4 have a medium impact, 3 have a high impact, 2 are deleterious, and 2 are probably damaging. In terms of PTM’s, there are 24 phosphorylation sites and 22 acetylation sites. Most of the common mutations mapped onto intron.

**MYH7:** Likewise, MYH6, it is a protein found as a component of the myosin protein in the heart, and having a mutation is associated with enlarged heart and other CVDs [34, 35]. A total of 753 mutations were mapped onto the MYH7 sequence (Figure 4X): 2 are missense mutations, 10 are splice mutations, 494 are intron mutations, 63 are silent mutations, 180 are 5' flank mutations, and 4 are 3' UTR mutations. Of these mutations, 1 has medium impact, 1 has high impact, 1 is tolerated, 1 is deleterious, and 2 are possibly damaging. In terms of PTM's, 111 are phosphorylation sites, 30 are acetylation sites, 4 are O-linked glycosylation sites, and there is 1 methylation site. The most numerous mutations mapped on the sequence are intron mutations followed by the 5' flank mutation.

**NOS3:** A protein primarily found in blood vessels that helps synthesize nitric oxide, and reduction in levels of NOS3 have been associated with HF [36]. We found a total of 1239 mutations mapped onto the NOS3 sequence (Figure 5A): 33 are missense mutations, 1 is a splice mutation, 922 are intron mutations, 53 are silent mutations, 225 are 5' flank mutations, 2 are 5' UTR mutations, and 3 are 5' UTR mutations. Of these mutations, 31 are tolerated, 1 is possibly damaging, and 1 is probably damaging. In terms of PTM's, there are a total of 23 phosphorylation sites, 2 acetylation sites, 1 O-linked glycosylation site, 13 are S-nitrosylation sites, 1 is a methylation site, and 2 are glutathionylation sites. Most of the mutations were intron mutations, followed by 5' flank mutations.

**NPPA:** Gene that generates atrial natriuretic peptide which functions to decrease blood pressure [37]. Mutations in NPPA are associated with HF, high blood pressure, atrial fibrillation, and heart attack. A total of 267 mutations were mapped onto the NPPA sequence (Figure 5B). There are 2 missense mutations, 8 are truncating mutations, 125 are intron mutations, 110 are 5' flank mutations, and 22 are 3' UTR mutations. Of these mutations, only the 2 missense mutations received a functional impact score and they both have low impact. There are no PTM's on the sequence. Most of the mutations are intron mutations, followed closely by 5' flank mutations.

**NPPB:** A hormone specific to the heart that functions to decrease blood pressure, and high levels are linked to HF [38]. We mapped a total of 524 mutations onto the NPPB sequence (Figure 5C). There is 1 splice mutation, 207 5' flank mutations, 289 3' flank mutations, 3 5' UTR mutations, and 24 3' UTR mutations. There are 2 phosphorylation sites and 7 O-linked glycosylation sites. Most of the mutations are 3' flank, followed closely by 5' flank mutations.

**NPPC:** A protein that can cause natriuresis, blood vessels to widen, and blood pressure to drop. NPPC levels tend to be high in HF patients [39]. A total of 283 mutations were mapped onto the NPPC sequence (Figure 5D). There are 8 intron mutations, 147 5' flank mutations, and 128 3’ flank mutations. In terms of PTM’s, there is only 1 acetylation site and 1 O-linked glycosylation. Most of the mutations were mapped onto 5’ flank followed by the 3’ flank mutation.

**NPR1:** A receptor protein that is found in many different organs including the heart, and mutations in NPR1 have been linked to HF, high blood pressure, and other CVDs [40]. We found 586 mutations mapped onto the NPR1 sequence (Figure 5E). There are 9 missense mutations, 204 intron mutations, 85 are 5’ flank mutations, 257 are 3’ flank mutations, 5 are 5’ UTR mutations, and 26 are 3’ UTR mutations. Of these mutations, 8 have neutral impact and 1 has low impact. In terms of PTM’s, there are 15 phosphorylation sites and 2 acetylation sites. Most of the mutations mapped onto the sequence are 3’ flank mutations followed by intron mutations.

**NR3C2:** A protein receptor expressed in many organs including the heart. Binding of aldosterone to NR3C2 in the heart has been associated with HF, high blood pressure, enlargement of the heart, and other CVDs [41]. A total of 18,645 mutations were mapped onto the NR3C2 sequence (Figure 5F). There are 35 missense mutations, 26 splice mutations, 18,278 intron mutations, 39 silent mutations, 126 5’ flank mutations, 67 3’ flank mutations, 3 5’ UTR mutations, and 71 3’ UTR mutations. Of these mutations, only the missense mutations received a functional impact score, and all 35 were rated benign by mutation assessor and tolerated by SIFT. In terms of PTM’s, there are 29 phosphorylation sites and 1 acetylation site. Most of the mutations mapped onto the sequence are intron mutations.

**PIK3C2A**: A gene that has multiple roles, and mutations and irregular expression have been associated with HF and other CVDs [42, 43]. A total of 3970 mutations were found on the PIK3C2A sequence (Figure 5G). There are 5 missenses, 3442 introns, 63 silent, 266 5’ flank, 127 3’ flank, 7 5’ UTR, and 60 3’ UTR mutations. Of these mutations, only the 5 missense mutations received a functional impact score and all 5 received a score of medium impact from mutations assessor, tolerated from SIFT, and benign from PolyPhen2. In terms of PTM’s, there are 50 phosphorylation sites, 9 acetylation sites, 15 ubiquitination sites, and 3 methylation sites. Most were mapped to introns, followed by 5’ and 3’ flanks.

**PLN:** It is a protein that can affect contraction of heart muscle, and its elevated activity can cause HF and other CVDs [44]. We mapped a total of 608 mutations onto the PLN sequence (Figure 5H). There are 535 intron mutations and 73 3’ UTR mutations. Most of the mutations were mapped to introns but no functional impact scores were retrieved for any of the mutations. In terms of PTM’s, there are only 2 phosphorylation sites.

**REN:** A hormone that regulates blood pressure, and its high levels are linked to HF and other CVDs [45, 46]. A total of 800 mutations were mapped onto the REN sequence (Figure 5I). There are 631 introns, 10 silent, 151 5' flank, and 8 5' UTR mutations. In terms of PTM's, there are 6 phosphorylation sites and 1 N-linked glycosylation site. Most of the mutations were mapped to introns, followed by 5' flanks, but no functional impact scores were retrieved for any of the mutations.

**TNF:** A cytokine that induces inflammation, and high amounts are linked to HF and other CVDs [47]. A total of 138 mutations were mapped onto the TNF sequence (Figure 5J): 1 missense mutation, 26 intron mutations, 107 5' flank mutations, and 4 3' UTR mutations. The 1 missense mutation received a functional impact score of low from mutation assessor. In terms of PTM's, there are 3 phosphorylation sites, 1 O-linked glycosylation site, and 2 myristoylation sites. Most of the mutations were in 5' flanks.

**UTS2:** A protein that narrows blood vessels, and high levels are linked to CVDs including HF [48]. We mapped a total of 1170 mutations onto the UTS2 sequence (Figure 5K). There are 11 missense mutations, 324 intron mutations, 2 silent mutations, 619 5' flank mutations, 210 3' flank mutations, and 4 3' UTR mutations. Of these mutations, 11 have low functional impact, 28 are tolerated, 10 are tolerated with low confidence, and 38 are benign. There are no PTM's on this sequence. Most of the mutations were mapped to 5' flanks followed by introns.

**VCL:** A protein that influences the cytoskeleton, and mutations that affect levels in plasma are linked with HF [49]. We found a total of 3518 mutations mapped onto the VCL sequence (Figure 5L). There are 3132 introns, 71 silent, 199 5' flank, and 116 are 3' UTR mutations. No functional impact scores were retrieved for any of these mutations. In terms of PTM's, there are 57 phosphorylation sites, 20 acetylation sites, 12 ubiquitination sites, 5 methylation sites, 3 S-nitrosylation sites, 13 are malonylation sites, 1 glutathionylation site, and 3 succinylation sites. Most of the mutations were mapped to introns, followed by 5' flanks and 3' UTRs.

### 2. Mutation analysis of other CVD genes

**ATP2A2:** A protein that controls calcium levels in cardiac muscle and, therefore, influences contraction of cardiac muscle [50]. Abnormal expression of ATP2A2 is linked to CVDs. We found a total of 208 mutations mapped onto the ATP2A2 sequence (Figure 6A). There are 1 splice, 172 introns, 3 silent, 5 5' flank, 16 3' flank, 3 5' UTR, and 8 3' UTR mutations. No functional impact scores were retrieved for any of these mutations. In terms of PTM's, there are 32 phosphorylation sites, 17 acetylation sites, 13 ubiquitination sites, 2 S-nitrosylation sites, 3 methylation sites, 3 malonylation sites, 9 glutathionylation sites, and 2 nitration sites.

**CALD1:** A protein that affects actin and interacts with myosin in smooth muscle [51]. Mutations in CALD1 is associated with CVDs [52]. We mapped a total of 925 mutations onto the CALD1 sequence (Figure 6B). There are 3 missenses, 867 introns, 1 silent, 44 5' flank, 2 5' UTR, and 8 3' UTR mutations. Of these mutations, 1 is neutral, 1 has a low impact, 1 has a medium impact, 1 is tolerated low confidence, 1 is deleterious low confidence, 1 is deleterious, 2 is benign, and 1 is possibly damaging. In terms of PTM's, there are 72 phosphorylation sites, 18 acetylation sites, 14 malonylation sites, and 2 sumoylation sites.

**CD34:** A gene with many functions, including generation of red blood cells. Low levels of CD34 are linked to multiple CVDs [53]. A total of 161 mutations were mapped on the CD34 sequence (Figure 6C). There are 2 missenses, 1 splice, 95 intron, 1 5' flank, 33 3' flank, and 29 3' UTR mutations. 2 mutations were given a score of low functional impact, 2 are tolerated, and 2 are benign. In terms of PTM's, there are 12 phosphorylation sites, 1 acetylation site, and N-linked glycosylation site.

**CD40LG:** A protein that binds to the CD40 receptor and causes inflammation. Increased amounts are linked to many CVDs [54]. We discovered a total of 51 mutations mapped on the CD40L sequence (Figure 6D). There are 26 introns, 1 silent, 14 5' flank, and 10 3' UTR mutations. In terms of PTM's, there are 2 phosphorylation sites and 1 N-linked glycosylation site.

**DDX41:** A protein that binds to nucleic acids and plays a role in stimulating interferon release [55]. Mutations in DDX41 are linked to CVDs, and to acute myeloid leukemia [56, 57]. We found and mapped a total of 19 mutations onto the DDX41 sequence (Figure 6E). There is 1 splice, 9 introns, 6 5' flank, and 3 3' UTR mutations. In terms of PTM's, there are 22 phosphorylation sites, 5 acetylation sites, 1 ubiquitination site, and 2 sumoylation sites. Most of the mutations were mapped to introns, followed by 5' flanks.

**ENO2:** A protein mainly found in neurons and involved in glycolysis [58]. Increased amounts of ENO2 are linked to low survival of CVDs, specifically cardiac arrest [59]. We mapped a total of 26 mutations onto the ENO2 sequence (Figure 6F). There are 23 introns, 1 5' UTR, and 2 3' UTR mutations. In terms of PTM'S, there are 12 phosphorylation sites, and 10 acetylation sites.

**FADD:** A protein involved in the process of cell death [60]. Blocking the activity of FADD has been proven to alleviate symptoms and improve outcomes in CVDs [60]. We found a total of 41 mutations mapped onto the FADD sequence (Figure 6G). There are 2 missenses, 7 are intron, 29 5' flank, 1 5' UTR, and 2 3' UTR mutations. Of these mutations, 2 have low functional impact, 1 is tolerated, 1 is benign, 1 is deleterious, and 1 is probably damaging. In terms of PTM's, there are 11 phosphorylation sites and 1 glutathionylation sites.

**FGF2:** A protein with many functions on cell development, growth, and process of developing new blood vessels [61]. High amounts of FGF2 are linked to CVDs such as abnormal amounts of fluid in the pericardium [61]. We found a total of 434 mutations in the FGF2 sequence (Figure 6H). There is 1 missense, 381 introns, 3 silent, 21 5' flank, and 28 3' UTR mutations. Of these mutations, only the 1 missense mutation received functional impact scores. It received a score of neutral impact from mutation assessor, deleterious low confidence from SIFT, and benign from PolyPhen-2. In terms of PTM's, there are 15 phosphorylation sites, 5 acetylation sites, 3 malonylation sites, and 1 sumoylation site.

**FGF23:** A protein that controls the levels of phosphate and 1,25-dihydroxyvitamin D. High amounts of FGF23 are linked to CVDs [61]. A total of 45 mutations were found in the FGF23 sequence (Figure 6I). There is 1 missense, 38 introns, and 6 3' UTR mutations. Of these mutations, the single missense mutation was the only mutation to have received functional impact scores. It received a score of neutral impact from mutation assessor, tolerated low confidence from SIFT, and benign from Polyphen-2. In terms of PTM's, there are 10 phosphorylation sites and 1 O-linked glycosylation site.

**FLNA:** A protein that has effects on cell structure and movement. Mutations in FLNA are linked to CVDs, specifically morphological abnormalities of the heart [62]. A total of 75 mutations were mapped onto the FLNA sequence (Figure 6J). There are 2 missenses, 56 introns, 5 silent, 8 3' flank, 1 5' UTR, and 3 3' UTR mutations. Of these mutations, 1 has low functional impact, 1 has medium impact, 1 is tolerated, 1 is deleterious, 1 is benign, and 1 is probably damaging. In terms of PTM's, there are 175 phosphorylation sites, 78 acetylation sites, 11 ubiquitination sites, 13 S-nitrosylation sites, 48 malonylation sites, 11 glutathionylation sites, 3 succinylation sites, and 1 sumoylation site.

**GJB6:** It is a gap junction protein, which facilitates movement of molecules [63]. Mutations in GJB6 are associated with difficulty with CVDs [63, 64]. We mapped a total of 167 mutations onto the GJB6 sequence (Figure 6K). There is 1 missense, 78 introns, 34 5' flank, 51 3' flank, 1 5' UTR, and 2 3' UTR mutations. Of these mutations, only the single missense mutation received functional impact scores: medium impact from mutation assessor, tolerated from SIFT, and probably damaging from PolyPhen-2. Most of the mutations were mapped to introns, followed by the 5' and 3' flanks.

**GLMN:** A protein that plays a role in the development of blood vessels, and mutations in GLMN are associated with CVDs, and issues with vessel development [65]. We found a total of 121 mutations mapped to the GLMN sequence (Figure 6L). There is 1 splice, 118 introns, 1 silent, and 1 3' UTR mutations. None of the mutations were assigned functional impact scores. In terms of PTM's, there are 6 phosphorylation sites, 3 acetylation sites, and 4 ubiquitination sites.

**HBA1:** It is a component of hemoglobin, which functions to carry oxygen throughout the body [66]. Abnormal levels of hemoglobin are associated with CVDs [67]. We mapped a total of 10 mutations onto the HBA1 sequence (Figure 6M): 8 of the mutations are 5' flank and 2 are 3' flank mutations. No functional impact scores were assigned to any of the 10 mutations. In terms of PTM's, there are 14 phosphorylation sites, 10 acetylation sites, 3 O-linked glycosylation sites, and 1 S-nitrosylation site. Most of the mutations were mapped to 5' flanks, followed by 3' flanks.

**KANTR:** It is an mRNA molecule that does not generate a protein [68]. Mutations in KANTR are linked to CVDs, and uncontrolled movements and spasms [68]. We found a total of 153 mutations mapped to the KANTR sequence (Figure 6N): 116 introns, 1 silent, 4 5' flank, 22 3' flank, 2 5' UTR, 6 3' UTR, and 2 RNA mutations.

**LEMD3:** It is a protein in the inner nuclear membrane that interacts with Smad 2 and 3 and has effects on signaling [69]. Mutations in LEMD3 are linked to CVDs and diseases that affect bone such as Buschke-Ollendorf syndrome [70]. We mapped a total of 317 mutations onto the LEMD3 sequence (Figure 6O). There are 1 missense, 1 splice, 265 introns, 2 silent, 22 5' flank, 22 3' flank, and 4 3' UTR mutations. Only the single missense mutation received functional impact scores: low functional impact from mutation assessor, deleterious from SIFT, and benign from PolpyPhen2. In terms of PTM's, there are 69 phosphorylation sites, 8 acetylation sites, 3 ubiquitination sites, 1 methylation site, and 4 malonylation sites.

**MB:** A protein that binds to nitric oxide and oxygen, and alleviates damage due to lack of oxygen to the heart [71]. Lack of MB is linked to CVDs, specifically buildup of lipids in the heart [71]. We found a total of 157 variants in MB sequence (Figure 6P). There is 1 splice, 77 introns, 2 silent, 33 5' flank, 41 3' flank, 1 5' UTR, and 2 3' UTR mutations. In terms of PTM's, there are 8 phosphorylation sites and 8 acetylation sites.

**PDPN:** A protein that has a role in the development of organs including the heart [72]. High amounts of PDPN are linked to CVDs, including heart attack [73]. We found a total of 246 mutations mapped to the PDPN sequence (Figure 6Q). There are 1 missense, 160 introns, 1 silent, 26 5' flank, 40 3' flank, 1 5' UTR, and 17 3' UTR mutations. Only the single missense mutation received functional impact scores: neutral by mutation assessor, tolerated by SIFT, and benign by PolyPhen-2.

**SLC2A1:** A gene that codes for GLUT1, which is a glucose transporter [74]. Irregular expression of SLC2A1 is associated with CVDs [75]. We mapped a total of 210 variants onto the SLC2A1 sequence (Figure 6R). There is 1 missense, 172 introns, 5 silent, 1 5' flank, 29 3' flank, 1 5' UTR, and 1 3' UTR mutation. Only the single missense mutation received a functional impact score. It was rated neutral by mutation assessor but was not given a score by SIFT and PolyPhen-2. In terms of PTM's, there are 11 phosphorylation sites, 3 acetylation sites, 2 ubiquitination sites, 1 N-linked glycosylation site, and 1 O-linked glycosylation site.

**SMUG1:** It generates a protein that removes uracil from DNA [76]. Low levels of SMUG1 are linked to CVDs and breast cancer [76]. A total of 104 mutations were mapped onto the SMUG1 sequence (Figure 6S). There are 1 missense, 1 truncating, 58 introns, 15 5' flank, 24 3' flank, 1 5' UTR, and 4 3' UTR mutations. Only the single missense mutation received a functional impact score. It received a score of neutral impact from mutation assessor and did not receive a score from either SIFT or PolyPhen-2.

**TAC1:** A gene that generates a protein called substance P [77]. Increased amounts of substance P are linked to CVDs, including inflammation of the heart [77]. We found a total of 92 mutations mapped onto the TAC1 sequence (Figure 6T). There are 32 introns, 20 5' flank, 35 3' flank, 5 are 3' UTR. In terms of PTM's, there are 2 methylation sites. Most mutations mapped to 3' flanks, followed by introns and 5' flanks.

**TEK:** It acts as a receptor protein for Angiopioetin-1 and has many functions including influencing the growth of blood vessels [78]. Mutations in TEK are linked to CVDs, especially abnormal formation of blood vessels and the heart [79]. A total of 1042 mutations were found in the TEK sequence (Figure 6U). There are 4 missenses, 938 introns, 8 silent, 24 5' flank, 60 3' flank, 1 5’ UTR, and 7 3’ UTR mutations. Of these mutations, 2 have neutral impact, 2 have low impact, 4 are tolerated, and 4 are benign. In terms of PTM’s, there are 16 phosphorylation sites and 2 N-linked glycosylation sites.

**TRPV1:** A protein channel whose activity is triggered by changes in temperature and pH [80]. TRPV1 activity is linked to CVDs, including high blood pressure, and thickening of the heart [81]. We found a total of 348 mutations mapped onto the TRPV1 sequence (Figure 6V). There are 8 missenses, 215 introns, 6 silent, 67 5’ flank, 46 3’ flank, 2 5’ UTR, and 4 3’ UTR mutations. Of these mutations, 5 have neutral impact, 2 have low impact, 1 has medium impact, 6 are tolerated, 2 are deleterious, and 8 are benign. In terms of PTM’s, there are 22 phosphorylation sites and 1 acetylation site.

**ZBTB8OS:** A protein involved in the process of RNA ligation via tRNA ligase [82]. ZBTB8OS can control the levels of XBP1, which are linked to number of CVDs and cancers such as B-cell leukemias [82]. A total of 181 mutations were mapped onto the ZBTB8OS sequence (Figure 6W). There is 1 splice, 150 introns, 1 silent, 23 3’ flank, and 6 3’ UTR mutations.

### Acknowledgements

We appreciate great support by the Pat and Jim Calhoun Cardiology Center, and Department of Genetics and Genome Sciences, at the UConn School of Medicine, UConn Health; Rutgers Institute for Health, Health Care Policy and Aging Research (IFH), and Rutgers Robert Wood Johnson Medical School (RWJMS), Rutgers Biomedical and Health Sciences (RBHS) at the Rutgers, The State University of New Jersey.

We would like to give special thanks to Dr. Christopher Bonin and Dr. Geneva Hargis for stylistic and native speaker corrections.

### Author contributions

ZA lead this study. ZA did RNA-seq data processing, quality checking, and downstream analysis. ZA developed MAV-clic and PROMIS-LCR, supervised JWES and GVViZ implementation, and performed cohort building and integrative clinical data analysis of consented patients. SZ provided bioinformatics expertise in variant data analysis and visualization. NP supported post-computational analysis and evaluation of results. ZA drafted the paper. All authors have participated in writing and review, and have approved paper for publication. BL proposed, supervised, and supported the study.

### Authors Bio

ZA is the Assistant Professor of Medicine – Tenure Track and Core Member at the Rutgers Institute for Health, Health Care Policy and Aging Research; and Department of Medicine – Division of General Internal Medicine, Rutgers Robert Wood Johnson Medical School, Rutgers Biomedical and Health Sciences, Rutgers University-New Brunswick. ZA is the Adjunct Assistant Professor at the Department of Genetics and Genome Sciences, UConn School of Medicine, UConn Health, CT; and Full Academic Member of the Rutgers Microbiology and Molecular Genetics; Center for Cancer Health Equity, Rutgers Cancer Institute of New Jersey; Rutgers Human Genetics Institute of New Jersey, NJ.

NP is the MD student at the Rutgers Robert Wood Johnson Medical School.

SZ is the Senior Scientist Associated with the Rutgers Cancer Institute of New Jersey.

BL is the Interim Chief Executive Officer, UConn Health; Executive Vice President for Health Affairs; Dean, UConn School of Medicine; Director, Pat and Jim Calhoun Cardiology Center; and Ray Neag Distinguished Professor of Cardiovascular Biology and Medicine. BL is an internationally recognized cardiovascular physician-scientist and national leader in academic medicine.
